# Analysis of gene expression and mutation data points on contribution of transcription to the mutagenesis by APOBEC enzymes

**DOI:** 10.1101/2021.04.14.439818

**Authors:** Almira Chervova, Bulat Fatykhov, Alexander Koblov, Evgeny Shvarov, Julia Preobrazhenskaya, Dmitry Vinogradov, Gennady V. Ponomarev, Mikhail S. Gelfand, Marat D. Kazanov

## Abstract

Since the discovery of the role of the APOBEC enzymes in human cancers, the mechanisms of this type of mutagenesis remain little understood. Theoretically, targeting of single-stranded DNA by the APOBEC enzymes could occur during cellular processes leading to the unwinding of DNA double-stranded structure. Some evidence points to the importance of replication in the APOBEC mutagenesis, while the role of transcription is still underexplored. Here, we analyzed gene expression and whole genome sequencing data from five types of human cancers with substantial APOBEC activity to estimate the involvement of transcription in the APOBEC mutagenesis and compare its impact with that of replication. Using the TCN motif as the mutation signature of the APOBEC enzymes, we observed a correlation of active APOBEC mutagenesis with gene expression, confirmed the increase of APOBEC-induced mutations in early-replicating regions, and estimated the relative impact of transcription and replication on the APOBEC mutagenesis, which turned out to be approximately equal in transcribed regions. We also found that the known effect of higher density of APOBEC-induced mutations on the lagging strand was highest in middle-replicating regions, and observed higher APOBEC mutation density on the sense strand, the latter bias positively correlated with the gene expression level.

**Bullet points:**

– The APOBEC mutagenesis rate is higher in actively expressed genes
– The APOBEC mutation density is higher on the sense strand
– The lagging/leading strand ratio of the APOBEC mutational density is highest in middle-replicating regions

## Introduction

Apolipoprotein B mRNA editing catalytic polypeptide-like (APOBEC) is a family of enzymes of the human innate immune system, whose known role is the defense against viruses and transposable elements (Salter et al. 2016). The APOBEC enzymes bind to single-stranded viral DNA and deaminate cytosine, leading to C > T and C > G substitutions in the TpC context (Shi et al. 2017). Recently, APOBEC enzymes have been implicated in cancer mutagenesis (Roberts et al. 2012; Nik-Zainal et al. 2012; Burns et al. 2013a) with APOBEC-associated mutations detected in many types of human cancer, including breast, lung, bladder, head/neck, and cervical cancers (Alexandrov et al. 2013; Burns et al. 2013b; Roberts et al. 2013). In a majority of these cancer genomes, the APOBEC-signature mutations were found clustered in DNA and located on the same DNA strand (Roberts et al. 2012; Nik-Zainal et al. 2012). In addition, cancer genomes enriched in APOBEC-induced mutations also contain mutations with the APOBEC signature that are not positionally clustered along the genome. As the APOBEC enzymes have a strong specificity toward single-stranded DNA (ssDNA), it has been suggested that the enzymes generate mutation during one or several cellular processes associated with the unwinding of double-stranded human DNA, such as DNA repair, replication, or transcription (Roberts and Gordenin 2014). However, the exact mechanisms of APOBEC-associated mutagenesis remain unknown (Petljak et al. 2019).

During DNA replication, ssDNA regions are transiently formed behind the replication fork and theoretically can serve as a substrate for the APOBEC enzymes. Furthermore, nucleotide polymerization on the lagging strand runs in the opposite direction and requires formation of ssDNA loops (Hamdan et al. 2009; Pandey et al. 2009). Indeed, recent papers indicate that the APOBEC mutagenesis is associated with replication, as the density of APOBEC-induced mutations has a strong bias toward the lagging replication strand (Seplyarskiy et al. 2016) and is relatively higher in early-replicated regions (Kazanov et al. 2015).

During transcription, RNA polymerase binds to the antisense strand of DNA, leaving the other, sense strand single-stranded and hence potentially exposed to the APOBEC mutagenesis. Additionally, formation of R-loops, triple-stranded nucleic acid structures comprised of synthetized RNA hybridized with the DNA antisense, and single-stranded sense DNA (Sollier and Cimprich 2015), may facilitate the APOBEC access to the transient ssDNA in the non-transcribed strand. The first evidence for the link between the APOBEC mutagenesis and transcription was obtained in whole-genome, exome, and transcriptome study of bladder cancer (Nordentoft et al. 2014) that demonstrated the correlation of APOBEC-signature mutation rate with the mean expression level, and the bias towards the sense strand. Recent study in yeasts demonstrated susceptibility of the sense strand of tRNA genes to APOBEC mutagenesis, which were mutated 1000-fold times more frequently than the non-tRNA genomic regions (Saini et al. 2017). On the other hand, a study analyzing the distribution of APOBEC-induced mutations across genomes of 119 breast and 24 lung cancer samples (Kazanov et al. 2015) did not find statistically significant difference of the density of APOBEC-induced mutations between transcribed and non-transcribed genomic regions, leaving the relevance of transcription to the APOBEC mutagenesis in question.

Here, we analyzed the whole genome and transcriptome sequencing data on 505 tumors across 14 cancer types (Fredriksson et al. 2014), in an attempt to study the connection between the APOBEC mutagenesis and transcription. Our results point on the important role of transcription in APOBEC mutagenesis. That includes higher mutation load in actively expressed genes and on sense strand, presumably driven by the facilitated access of APOBEC enzymes to the single-stranded sense strand during the process of transcription.

## Results

### Selection of the APOBEC mutational signature for the analysis of human cancer genomes

To analyze the distribution of APOBEC-induced mutations along the genome and its connection with the replication and transcription one need to distinguish single-base substitutions (SBS) presumably generated by the APOBEC mutagenesis from all other SBS. Such classification of mutational data can be done using the mutational signature attributed to the APOBEC enzymes. Previous studies suggested to use TCW (W stands for A and T) (Roberts et al. 2013) or TCN (Burns et al. 2013b) motifs as the APOBEC mutational signature. Previously, using the TCW mutational signature, we observed the increased density of APOBEC-induced mutations in early-replicating regions (Kazanov et al. 2015) supplemented by a small shift in the same direction for the distribution of mutations not conforming to the TCW motif. This observation allowed us to speculate that APOBEC enzymes substantially target DNA outside of the TCW motif in human cancers and to use the TCN motif as more appropriate in this case. To validate this approach to the considered dataset, we calculated the distributions of the SBS density along the replication timing, while grouping single-base substitutions by their three-nucleotide contexts, i.e. considering all possible 5’ and 3’ bases of the mutated nucleotide (Supplemental File 1). We considered five cancer types having substantial numbers of samples enriched with APOBEC mutagenesis (see Methods; Fig.1a), namely, breast carcinoma (BRCA, 96 samples), bladder carcinoma (BLCA, 21 samples), head and neck carcinoma (HNSC, 27 samples), lung adenocarcinoma (LUAD, 46 samples), and lung squamous cell carcinoma (LUSC, 45 samples) (Fredriksson et al. 2014). As expected, the slope of the density distribution of APOBEC-induced SBS along the replication timing was negative for the TCA and TCT triplets (mutation density decreased toward the late-replicated regions, Fig. 1c) and, noticeably, also for the TCC and TCG triplets, as demonstrated for representative samples of five cancer types in Figure 1d. The slopes of the density distribution for all other triplets were mainly positive (Fig. 1b). In some cases (Fig. 1d, LUSC), slopes of the density distribution for TCN triplets were also positive, but still sufficiently smaller than the slopes for other triplets. This apparently reflects the mixed origin of mutagenesis in particular triplets, as each TCN motif contains mixture of mutations generated by different types of mutagenesis, i.e. not only SBSs induced by APOBEC-mutagenesis, but also SBSs generated from other sources. This can offset the effect of higher density of APOBEC-induced mutations in early replication regions, as the mutation density of most cancer signatures is relatively higher in late replication regions (Koren et al. 2012; Donley and Thayer 2013; Lawrence et al. 2013; Liu et al. 2013; Polak et al. 2014; Sima and Gilbert 2014; Schuster-Bockler and Lehner 2012). We argue that similar effects of higher mutation density in early-replicated regions both in TCW and TCS motifs, as well as known TCN specificity of the APOBEC enzymes toward viral DNA, demonstrate that TCC and TCG triplets are also targeted by APOBEC mutagenesis in human cancers, hence supporting our use of a less stringent TCN motif as the APOBEC mutational signature.

**Figure 1.**
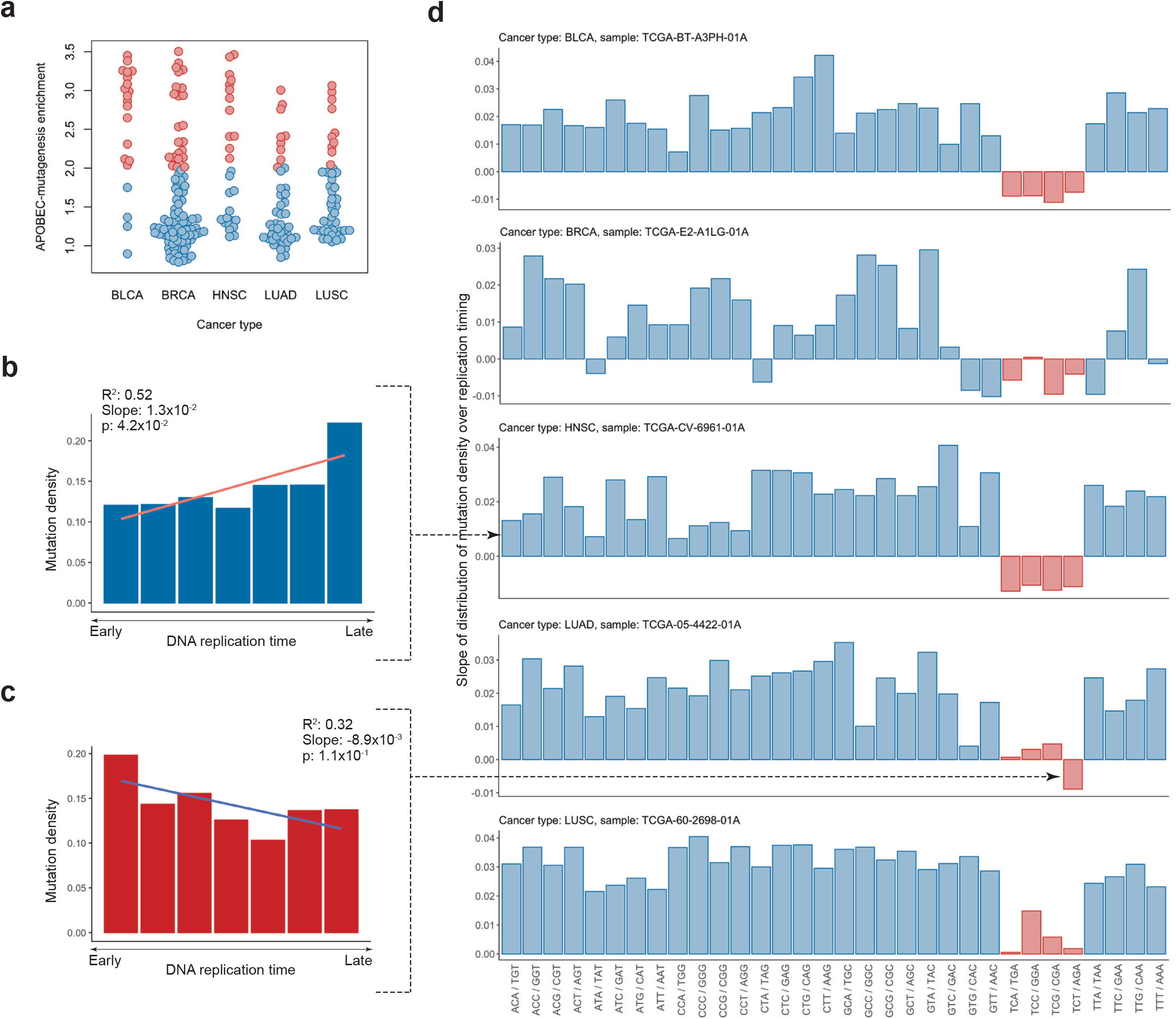
(a) Activity of APOBEC mutagenesis in samples from five cancer types. Samples enriched with APOBEC mutagenesis (empirical threshold on enrichment = 2.0) are highlighted in red. (b, c) Examples of the mutational density distribution over replication timing with positive and negative slopes for the ACA/TGT motif of TCGA-CV-6961-01A sample of the HNSC cancer and the TCT/AGA motif of TCGA-05-4422-01A sample of the LUAD cancer, respectively. (d) Representative cancer samples with the calculated slopes of the mutation density distribution over replication timing for all trinucleotide motifs with the substitution in the central nucleotide. Trinucleotides TCN are highlighted in red.

### The effect of the high density of APOBEC-induced mutations in early-replicated regions is more prominent in the case of TCN mutational signature

Using the newly defined APOBEC mutational signature (TCN motif) we attempt to confirm the earlier observed effect of the increased density of APOBEC-induced SBSs in early-replicated regions in human cancers (Kazanov et al. 2015). We calculated the slopes of distributions of the relative APOBEC mutation density over replication timing (see Methods). The results for five cancer types are shown in Figure 2. In plots for all cancer types, the slopes of the relative mutation density decrease with the increase of the APOBEC-enrichment of a sample (the APOBEC enrichment is a proxy for the activity of APOBEC enzymes in a particular sample, see Methods). This means that the APOBEC mutagenesis rate correlates with the increased density of APOBEC-induced mutations in early-replicating regions. The most profound effects can be seen for BLCA (slope of the regression line, *k*=–8.02×10^−3^, *p*-value =9.92×10^−3^), LUAD (*k*=– 1.15×10^−2^, *p*=3.44×10^−7^) and LUSC (*k*=–1.07×10^−2^, *p*=2.21×10^−9^) cancers, moderate effect for HNSC (*k*=–6.61×10^−3^, *p*=6.04×10^−3^), and less prominent effect for BRCA (*k*=–2.08×10^−3^, *p*=1.36×10^−2^). This effect was not so prominent when only the TCW motif without TCS triplets is used as the APOBEC mutational signature (Figure S1) as all slopes of trend lines were higher or equal than those for the TCN motif: BLCA *k*=–4.23×10^−3^, p=1.36×10^−1^; BRCA *k*=–1.17×10^−4^, *p*=1.36×10^−1^; HNSC *k*=–4.49×10^−3^, *p*=4.28×10^−2^; LUAD *k*=–1.19×10^−2^, *p*=1.64×10^−6^; LUSC *k*=– 1.03×10^−2^, p = 1.19×10^−7^.

**Figure 2.**
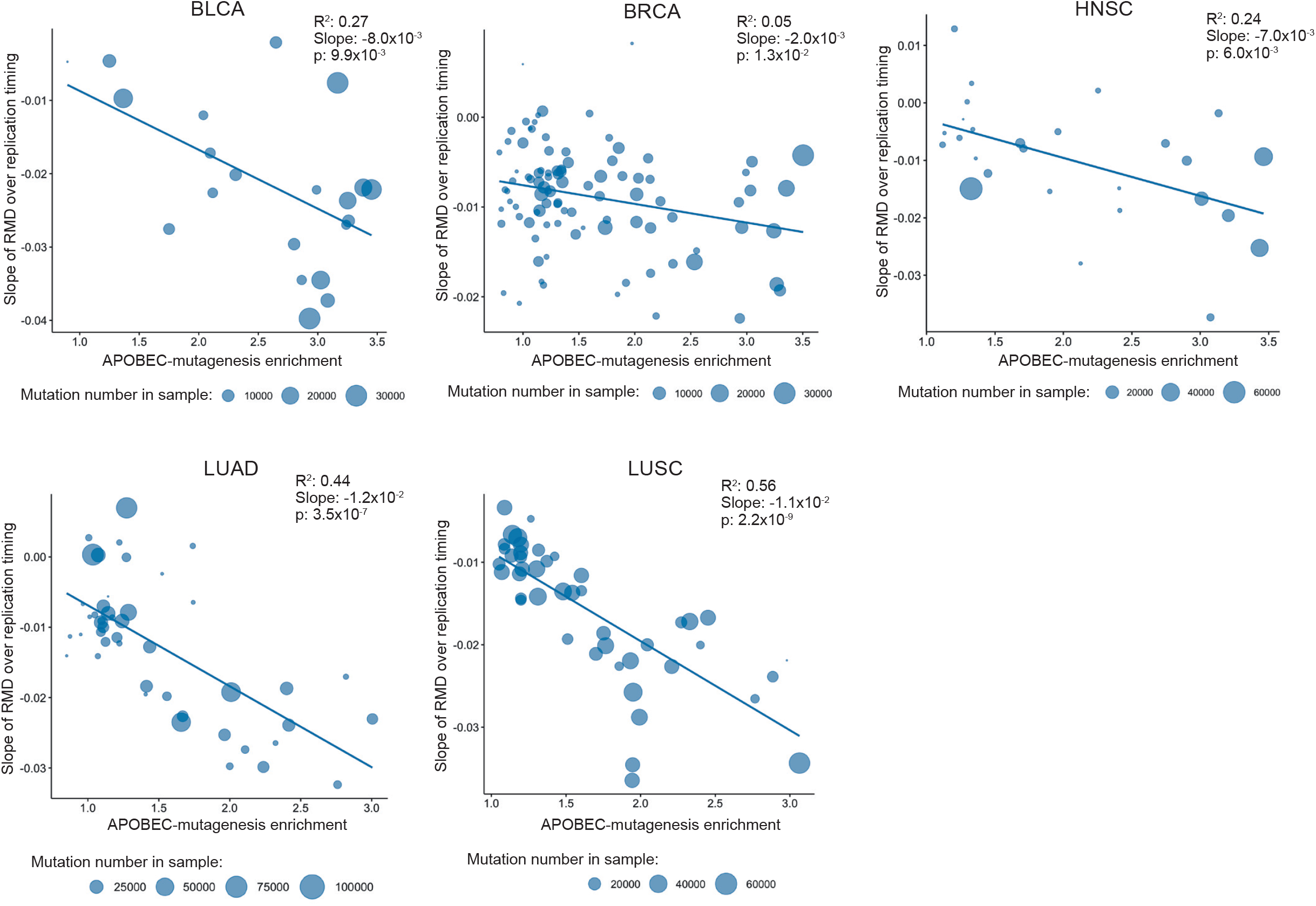
The slopes of the relative mutational density (RMD) distribution (see Methods) of APOBEC-induced SBSs (TCN motif) over replication timing as dependent on the activity of APOBEC mutagenesis for samples from five cancer types. The vertical coordinate is the estimated slope of the APOBEC-induced RMD over replication timing as shown in Supplemental file 1 or Figures 1b and 1c.

### APOBEC-mutagenesis is associated with higher mutation density in actively expressed genes

To elucidate possible relationship between transcription and APOBEC mutagenesis, we analyzed gene expression data associated with the studied cancer samples. We estimated the distribution of APOBEC-induced mutations in groups of genes stratified by expression levels. For each cancer sample we divided all genes into seven expression groups (bins) (see Table S4) and calculated the mutational density for each bin. Similar to the replication timing analysis, for each bin we calculated the relative APOBEC mutation density as the difference between the density of APOBEC-induced SBSs and other SBSs in cytosines in genome regions associated with the given expression bin. The results for five cancer types are presented in Figure 3. To check that the distribution of mutational density of non-APOBEC-induced SBSs in cytosines is not substantially different from the distributions of SBSs in other nucleotides, we calculated the distributions of mutational density over gene expression levels for all triplets (Supplemental File 2).

**Figure 3.**
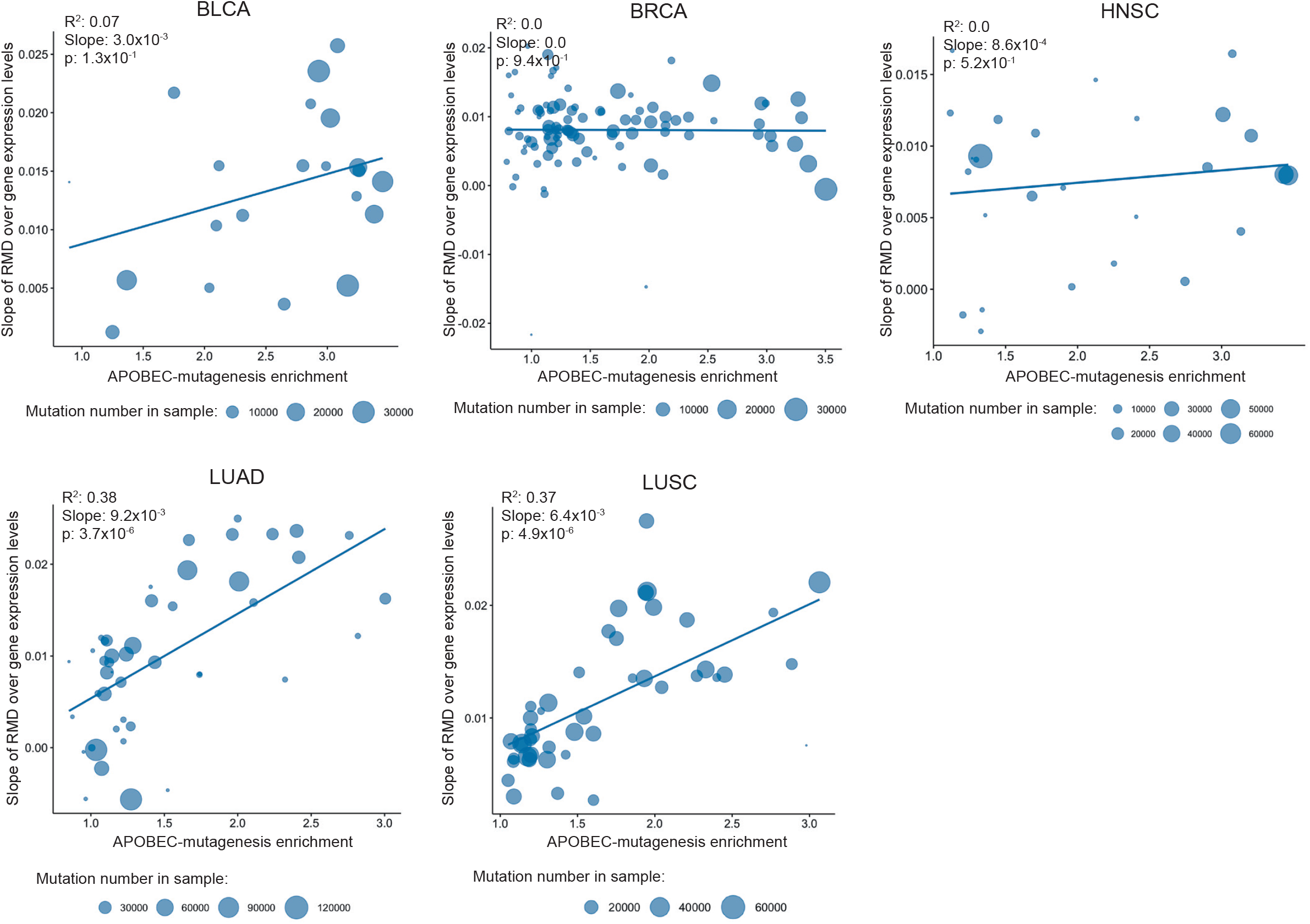
The slopes of the relative mutational density (RMD) distribution of APOBEC-induced SBSs (TCN motif) over gene expression levels as dependent on the activity of APOBEC mutagenesis for samples from five cancer types. The vertical coordinate is the estimated slope of the APOBEC-induced RMD over gene expression levels as shown in Supplemental file 2.

Figure 3 shows that stronger APOBEC signature enrichment of the sample corresponds to a steeper slope of the relative mutation density over gene expression levels, i.e. the activity of APOBEC mutagenesis is associated with the increased density of APOBEC-induced mutations specifically in actively expressed genes. This effect is strong for LUAD (slope of the regression line, *k*=9.22×10^−3^, *p*-value=3.73×10^−6^) and LUSC (*k*=6.4×10^−3^, *p*=4.92×10^−6^), noticeable for BLCA (*k*=3.0×10^−3^, *p*=1.3×10^−1^), weak for HNSC (*k*=8.63×10^−4^, *p*=5.24×10^−1^), and not visible for BRCA (*k*=–5.96×10^−5^, *p*=9.4×10^−1^). This effect is also prominent when the TCW motif is used as the APOBEC mutational signature (Figure S2).

### Lagging/leading strand ratio of APOBEC-induced mutational density is maximal at the middle of replicating timing

Further, we investigated how the known effect of high density of APOBEC-induced SBSs on the lagging strand (Haradhvala et al. 2016; Seplyarskiy et al. 2016) relates to the replication timing. Firstly, we confirmed the general effect of increased APOBEC-induced mutational density on the lagging strand by calculating the lagging/leading mutational density ratio for APOBEC-enriched samples (APOBEC enrichment > 2.0) from the considered dataset. We compared the results with ratios calculated for samples with low APOBEC activity (APOBEC enrichment < 2.0). As a control, we considered mutations in cytosines not conforming to the TCN motif in low APOBEC-enriched samples, so as to decrease as much as possible the influence of the APOBEC mutagenesis. Indeed, both low APOBEC-enrichment value of a sample and the mutation triplet not conforming to the APOBEC signature should decrease the probability that mutations taken as a control are APOBEC-induced. The mean lagging/leading mutational density ratio of APOBEC-induced SBSs in APOBEC-enriched samples was 1.35 against 1.0 for SBSs in cytosines excluding the TCN motif of low APOBEC activity samples (Mann–Whitney–Wilcoxon *p*-value=3.4×10^−4^) for BLCA, 1.41 vs. 1.05 for BRCA (*p*=3.3×10^−13^), 1.29 vs. 1.03 for HNSC (*p*=1.2×10^−7^), 1.24 vs. 1.0 for LUAD (*p*=2.2×10^−8^) and 1.3 vs. 0.99 for LUSC (*p*=6.3×10^−10^), respectively.

Then, we measured the lagging/leading strand ratio of APOBEC-induced mutational density along the replication timing (Figure 4). A combination of two known effects, increased density of APOBEC-induced mutations in early-replicating regions and on the lagging strand should yield the highest value of the lagging/leading strand ratio of the APOBEC-induced mutation density in the earliest replication timing bin. However, while in general the lagging/leading strand density ratio decreased from the early to late replication time, surprisingly, the highest values of this ratio were observed in the middle of the replication timing. Thus, the mean value of the lagging/leading strand ratio of APOBEC-induced SBS density over the samples was maximum at the third bin (numbered from early to late replication time) for all types of cancer: BLCA 1.5, BRCA 1.55, HNSC 1.43, LUAD 1.35, and LUSC 1.41. To estimate the statistical significance of this observation, we repeatedly randomly shuffled mutations between the lagging and leading strands (see Methods). The calculated *p*-values (BLCA: *p*=4.8×10^−12^, BRCA: *p*=6.0×10^−12^, HNSC: *p*=1.2×10^−8^, LUAD: *p*=1.8×10^−3^, LUSC: *p*=5.6×10^−6^) indicate that the observed effect is statistically significant (Figure S10). As a control, we observed that the lagging/leading strand ratio of mutational density over replication timing for other SBSs in cytosines was relatively flat (Figure S3). To make sure that the distribution in cytosines not conforming to the APOBEC signature is an appropriate representation of the background mutagenesis, we calculated the distributions of lagging/leading strand mutational density ratio in all triplets (Supplemental File 3) and confirmed it by a manual inspection.

**Figure 4.**
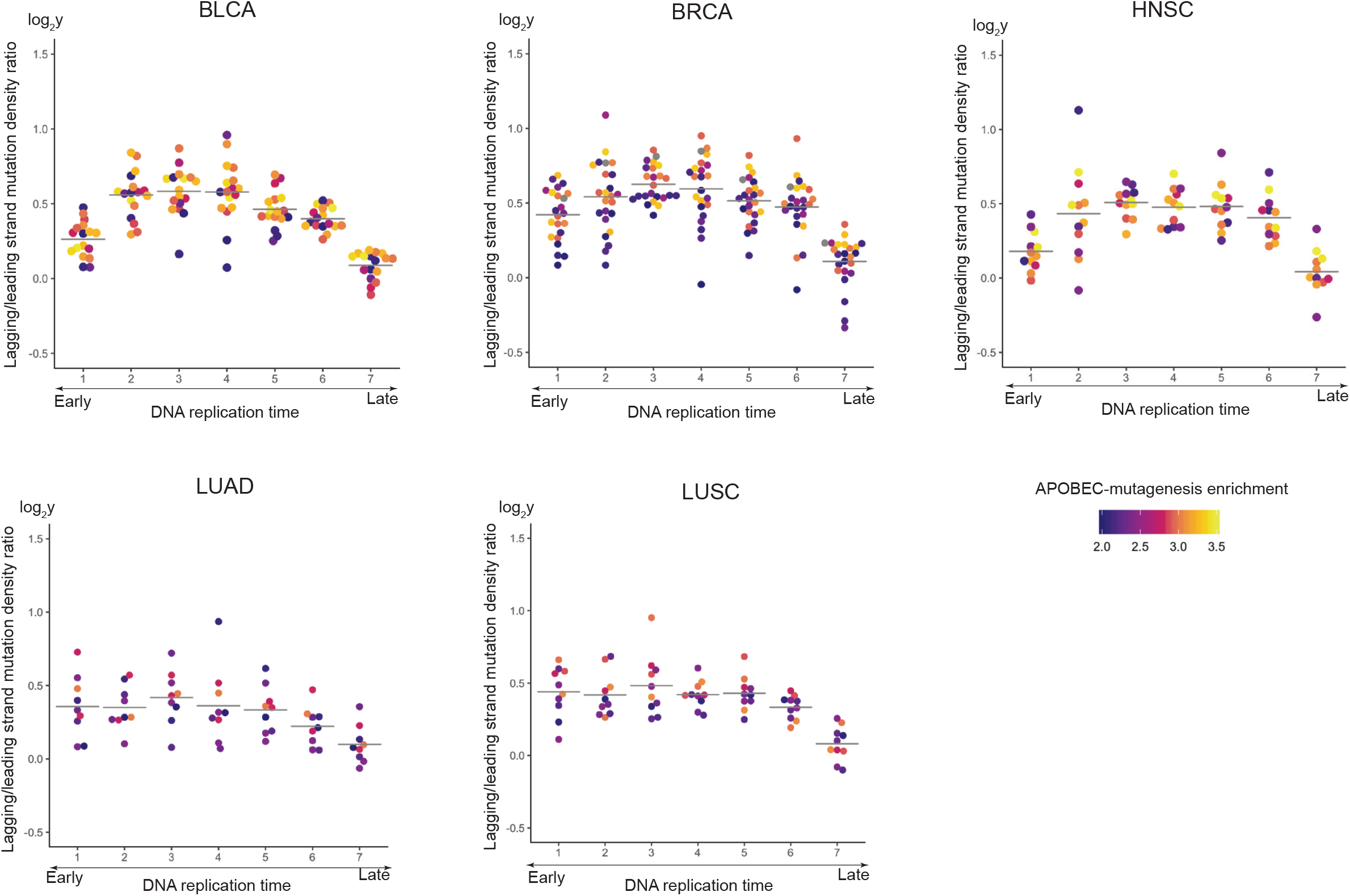
Dependence of the lagging/leading strand ratio of APOBEC-induced SBS density on the replication timing for samples from five cancer types. The horizontal lines show the average lagging/leading strand ratio values.

### The prevalence of APOBEC-induced mutations on the sense strand versus the antisense strand positively correlates with the gene expression level

Then, we compared APOBEC mutagenesis between the sense and antisense strands during transcription. We found a statistically significant increase of the APOBEC-induced SBS density on the sense strand, whereas for other SBS in cytosines we observed increased mutational density on the antisense strand. The mean sense/antisense strand density ratio of APOBEC-induced SBSs in APOBEC-enriched samples was 1.13 as compared with 0.74 for SBS in cytosines of low APOBEC activity samples (the Mann–Whitney–Wilcoxon test *p*-value=6×10^−3^) for BLCA, 1.04 vs. 0.98 for BRCA (*p*=8.2×10^−2^), 1.03 vs. 0.87 for HNSC (*p*=1.8×10^−2^), 1.1 vs. 0.65 for LUAD (*p*=1.3×10^−4^), and 1.08 vs. 0.69 for LUSC (*p*=6.3×10^−10^), respectively.

We also calculated the sense/antisense strand ratio of the APOBEC-induced mutational density over groups of genes stratified by expression levels. Figures 5, S8a shows that the sense/antisense strand ratio of APOBEC-induced SBS density increases with the gene expression level. This effect is observed for all considered cancer types: BLCA (slope of the regression line, *k*=7.7×10^−2^, *p*-value=1.8×10^−13^), BRCA (*k*=4.7×10^−2^, *p*=1.27×10^−6^), HNSC (*k*=3.65×10^−2^, *p*=2.02×10^−2^), LUAD (*k*=4.21×10^−2^, *p*=4.82×10^−4^), and LUSC (*k*=6.74×10^−2^, *p*=6.63×10^−9^). We suggest that this effect is associated both with the availability of the sense strand exposed in the single-stranded conformation for targeting by APOBEC enzymes (Jinks-Robertson and Bhagwat 2014) and with the repairing of the targeted cytosines on the antisense strand by the transcription-coupling repair (TCR) (Fousteri and Mullenders 2008; Hanawalt and Spivak 2008; Spivak and Ganesan 2014). Contrary to this tendency, the sense/antisense strand ratio for other SBS in cytosines (Figures S4, S8b, all triplets – Supplemental File 4) decreased with the increasing of gene expression level: BLCA (*k*=–1.5×10^−1^, *p*=7.83×10^−3^), BRCA (*k*=–2.47×10^−3^, *p*=8.09×10^−3^), HNSC (*k*=–8.66×10^−2^, *p*=5.57×10^−3^), LUAD (*k*=–1.37×10^−1^, *p*=1.48×10^−12^), and LUSC (*k*=–9.83×10^−2^, *p*=2.74×10^−23^). The latter effect might be associated with the smoking-based mutagenesis targeting guanines. Indeed, it is known that in smoking-related tumor genomes, guanine substitutions occur more frequently on the sense strand due to the transcription-coupled repair of the targeted guanines on the antisense strand (Pleasance et al. 2010), hence, reducing the density of the SBS in (complementary) cytosines on the sense strand.

**Figure 5.**
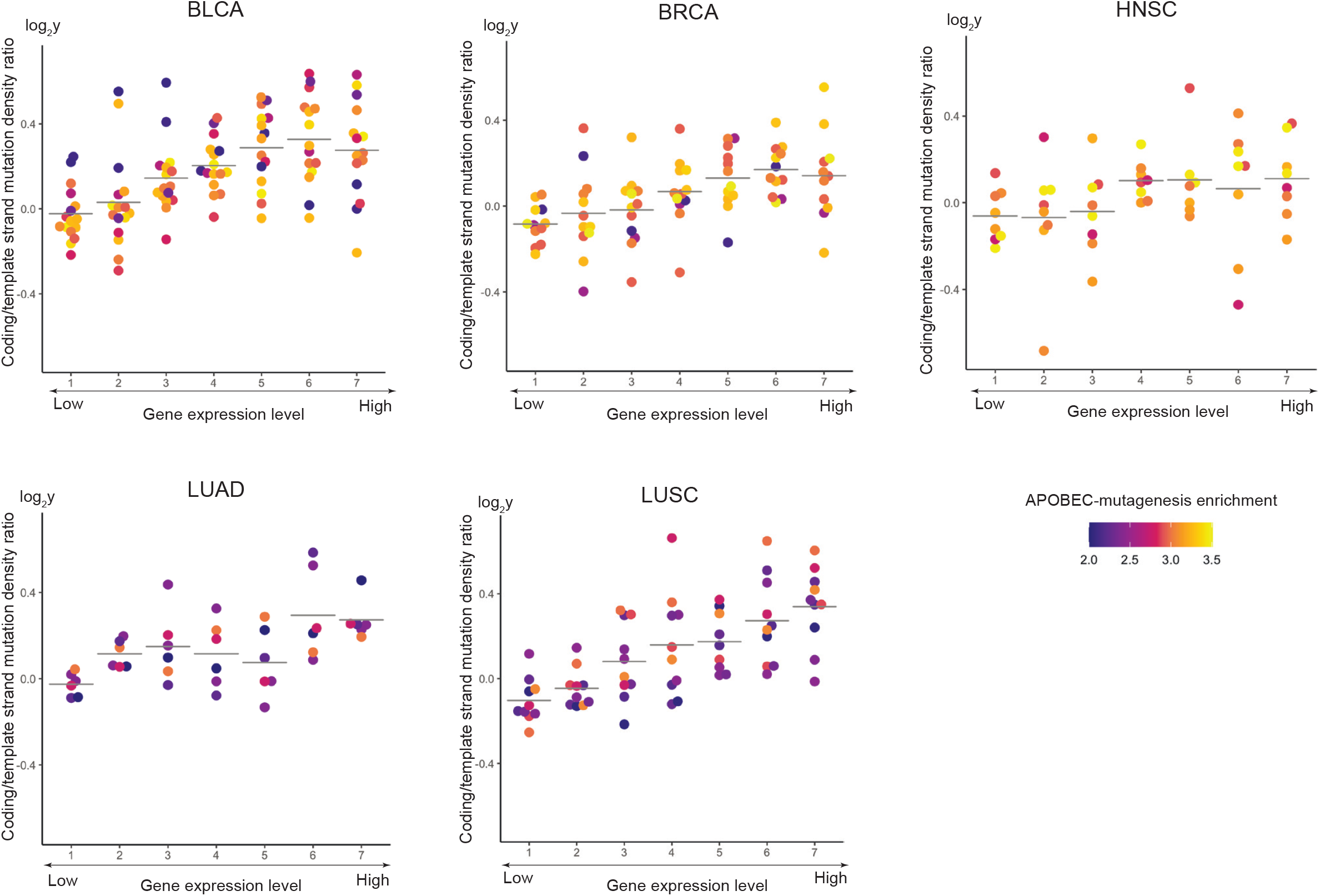
Dependence of the sense/antisense strand ratio of APOBEC-induced SBS density on the gene expression level for samples from five cancer types. The horizontal lines show the average sense/antisense strand values.

We also analyzed whether the density of APOBEC-induced mutations in gene regions depends on the mutual direction of replication and transcription. The comparison of genes with co-direction of replication and transmission and genes with anti-direction of these processes did not yield a statistically significant difference between the dependences of APOBEC-induced mutational density on the gene expression level in these two cases(Figure S9).

### Both replication timing and transcription contribute to the mutagenesis by APOBEC enzymes

Then, we analyzed whether both replication timing and gene expression influence APOBEC mutagenesis or only one feature is causative and other one is just correlated with the former. Firstly, we calculated the ratios of the transcribed to intergenic number and density of APOBEC-induced SBSs (Figures 6a, Figure S5). This showed that when the level of APOBEC mutagenesis increased, the total number and density of APOBEC-induced SBSs in gene regions also increased, in comparison with the total number and density of APOBEC-induced SBSs in intergenic regions. For samples with the strongest APOBEC-enrichment, the total number and density of mutations in transcribed regions increased almost to the level of the corresponding values for intergenic regions. However, as gene-dense regions of the genome is associated with early-replication domains (Woodfine et al. 2004), the increase of APOBEC-induced mutations in genes can be associated both with transcription and replication.

**Figure 6.**
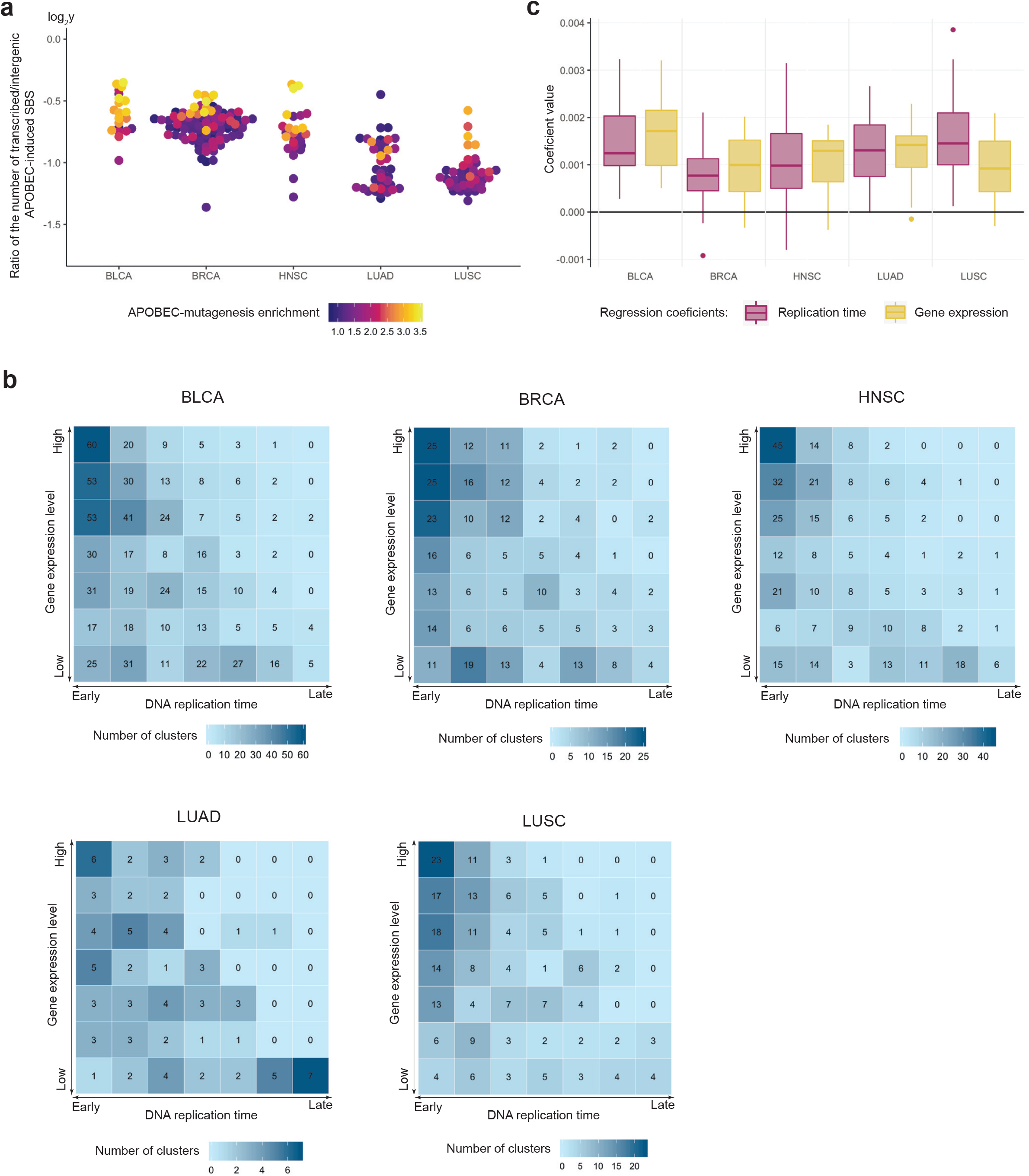
(a) Ratio of the number of APOBEC-induced mutations in gene/intergenic regions of samples from five cancer types. (b) Distribution of APOBEC-induced mutation clusters over replication timing and gene expression. (c) Regression coefficients reflecting the impact of replication timing and gene expression generated by the linear model (see Methods) approximating the density of APOBEC-induced SBS in gene regions.

To clarify the interdependence of transcription and replication in APOBEC mutagenesis, we calculated the number of SBS clusters over both replication timing and gene expression. Figure 6b shows that, for a particular replication timing bin, the number of APOBEC-induced SBS clusters grows with increasing expression level. The number of APOBEC-induced SBS clusters reaches maximum in regions corresponding to the highest gene expression level of the earliest replication timing bin. Thus, we can conclude that both replication and transcription contribute to the APOBEC mutagenesis. At the opposite, the maximum number of non-APOBEC-induced SBS clusters is concentrated in genome regions corresponding to the lowest expression level of the latest replication bin (Figure S6). We also present figures featuring the cluster density instead of the number of clusters by normalizing the numbers of clusters by the sizes of the respective genome regions (Figure S7). The observed trends remain the same after cluster normalization.

Then, we estimated the relative impact of the replication timing and gene expression on the APOBEC mutagenesis. For each sample we fit a linear model (see Methods) with two independent variables – replication timing and gene expression – and estimated their regression coefficients, reflecting the impact of each genomic feature. The coefficients’ absolute values for samples from five cancer types are shown in Figure 6c. As can be seen, the coefficient values of these two features are very close, so it cannot be concluded that one has stronger impact than the other in transcribed regions. To validate this conclusion, we also analyzed these data using two additional methods, LMG (Lindeman et al. 1980) and Analysis of Variance (ANOVA). Both methods have confirmed our initial conclusion that the contributions of the replication timing and gene expression both are significant, and their relative impacts are approximately equal (Figures S17, S18), although the results of ANOVA have shown that the relative impact of replication timing was higher in a larger number of samples. Despite the suggested approximately equal impact of replication and transcription, the total number of APOBEC-induced SBSs due to replication, taking into account mutations in intergenic regions, should be considerably larger than the number of SBSs due to transcription.

### Validation on PCAWG dataset

To validate our findings, we repeated the same analysis on a dataset available from the Pan-Cancer Analysis of Whole Genomes (PCAWG) project (Campbell et al. 2020). The PCAWG study is a project for identification of somatic and germline variations in both coding and non-coding regions of more than 2,600 cancer whole genomes across 38 cancer types. Similar to our initial analysis, we have selected cancer types with a substantial number of samples enriched with the APOBEC signature. Six cancer types were selected; five types as in the previous dataset, and the cervical cancer (CESC). It should be noted that most cancer samples of the (Fredriksson et al. 2014) dataset are also parts of the PCAWG dataset, but processed with a different mutation calling procedure. Thus, using the PCAWG dataset, we validated the results on both new cancer samples and types, and on the same samples with an alternative calling procedure. As shown in figures S11-S16, all findings – the increase of APOBEC-induced mutational density in highly expressed genes (Figure S12), the peak of elevated density of APOBEC-induced mutations on the lagging strand at the middle of the replication timing (Figure S13), the increased APOBEC-induced mutational density on the sense strand (Figure S14), and the approximately equal impact of replication and transcription on the APOBEC mutagenesis (Figures S15 and S16) – have been confirmed on PCAWG dataset.

## Discussion

While the implication of APOBEC enzymes in human cancer has been discovered eight years ago (Nik-Zainal et al. 2012; Roberts et al. 2012; Burns et al. 2013a), the mechanisms of APOBEC mutagenesis still are not understood well. The natural suspects are cellular processes associated with temporary unwinding of the DNA into the single-strand state, in particular, replication (Seplyarskiy et al. 2016; Kazanov et al. 2015) and, possibly, transcription (Taylor et al. 2014; Lada et al. 2015). Here, we have attempted to disentangle their contribution using whole-genome sequencing and gene expression data for cancers with substantial activity of APOBEC enzymes.

Accounting for the elevated density of APOBEC-induced mutations in early-replicating regions (Kazanov et al. 2015), we found indirect but strong evidence that the conventional mutational signature of APOBEC enzymes in human cancers, TCW, can be extended to TCN, as the TCC and TCG triplets also seem to be targeted by APOBEC. Using this mutational signature, we confirmed the higher density of APOBEC-induced mutations in early-replicating regions and found a strong correlation between the density of these mutations in genes and the level of gene expression.

The results for APOBEC-induced mutational clusters demonstrate stronger effect for bladder, head/neck, and breast cancer in comparison with lung cancers. The reason for the stronger effect for lung cancers in isolated (not clustered) mutations may be a better estimate of the background mutagenesis for lung cancers due to the higher number of mutations. As described in the Method section, we considered a mixture of APOBEC-induced and background mutagenesis in the TCN motif. It is possible that lung samples with a higher number of mutations allows us to better estimate the level of background mutagenesis and thus to estimate the proportion of APOBEC-induced mutations in more correct way than for samples with lesser number of mutations.

To estimate the relative impact of replication and transcription, we calculated the number of mutation clusters as a function of the replication timing and gene expression, and applied regression analysis to model the number of single-base substitutions. We conclude that both processes influence the activity of APOBEC mutagenesis with approximately equal impact in transcribed regions. The density of APOBEC-induced SBSs is almost equal in intergenic and transcribed regions for samples with the highest activity of APOBEC enzymes, meaning that the fraction of transcriptionally induced APOBEC mutations may be the same as the fraction of transcribed regions in the human genome, that is, about one fourth according to the GENCODE annotation (Frankish et al. 2019).

We have also analyzed possible strand asymmetry of APOBEC-induced single-base substitutions both for replication and transcription, and how their density changes over the replication timing and gene expression, respectively. We confirmed higher density of APOBEC-induced mutations on the lagging strand and found an unexpected distribution of the lagging/leading strand ratio of the mutational density over the replication timing, which reaches its maximum at the middle-replicating genome regions. This is not the case for other single-base substitution in cytosines, whose distribution between the replication strands is relatively uniform and independent from the replication timing. We speculate that this effect may be directly linked to the chromatin organization. Indeed, the middle of the replication timing is known for a dramatic switch from replication of euchromatin regions to replication of heterochromatin regions (Rhind and Gilbert 2013).

As for the transcriptional asymmetry, the density of APOBEC-induced SBS on the sense strand increases with the gene expression level, while the opposite is observed for other SBS in cytosines. The latter asymmetry could be due to the known smoking-associated damage of guanines and their repair on the antisense strand by transcription-coupled repair. This would lead to the prevalence of guanine substitutions on the sense strand, which is equivalent to the accumulation of cytosines substitutions on the antisense strand. We speculate that in APOBEC-enriched samples this asymmetry is compensated for and switched to the sense-strand cytosine-rich SBS asymmetry due to stronger action of APOBEC enzymes on heavily transcribed genes.

A mechanistic explanation for that might be that the sense strand is exposed during transcription in the single-stranded state and hence can be targeted by the APOBEC enzymes, whereas the antisense strand is occupied by the RNA polymerase complex (Jinks-Robertson and Bhagwat 2014). Thus, this mutational process, in addition to the transcription-coupled repair of cytosines on the antisense strand, could make the sense-strand cytosine-rich SBS asymmetry associated with the APOBEC mutagenesis stronger than smoking-associated sense-strand guanine-rich SBS asymmetry.

Overall, we have demonstrated an important role of transcription in mutagenesis by APOBEC enzymes in human cancer. Some of our observations, such as the increased density of APOBEC-induced SBS in the sense strand, have simple mechanistic explanations, while others, such as the fact that the lagging strand-associated bias in the density of APOBEC-induced mutations peaks in the middle-replicating regions, remain without underlying molecular mechanisms.

## Methods

### Dataset

Somatic alternations in 12 types of human cancer were taken from (Fredriksson et al. 2014). Indels were filtered out. Five cancer types, BLCA, BRCA, HNSC, LUAD, LUSC, which contained samples enriched with the APOBEC-mutagenesis signature (APOBEC-mutagenesis enrichment > 2.0, calculated as in (Roberts et al. 2013)), were selected for further analysis. Human genome assembly GRCh37/hg19 was used. Calculations were performed in InterSystems IRIS and MATLAB environments. Processing of computation-intensive subtasks was written in C++ and performed on computational cluster.

### Replication timing analysis

Replication timing data for MCF-7, IMR90, and NHEK cell lines was taken from the ENCODE database (Davis et al. 2018). Replication timing values were divided into seven intervals to create bins containing approximately equal number of the TCN motifs in each bin (Tables S1-S3). The mutation density D_APOBEC_ of the APOBEC mutagenesis in the genome regions corresponding to a particular replication timing bin was calculated as the number of single-base substitutions <u>C</u>->T or <u>C</u>->G in the T<u>C</u>N motif divided by the total number of the TCN triplets in these regions: *D*_APOBEC_ = *N*_APOBEC_ / *N*_TCN_. The relative mutation density of the APOBEC mutagenesis was calculated as *RD*_*APOBEC*_ = *D*_APOBEC_–*D*_NCN_, where *D*_NCN_ is the density of other single-base substitutions in cytosines. The replication data for the IMR90 cell line was used for analysis of the LUAD and LUSC mutational data; the NHEK cell line data, for HNSC and BLCA, and the MCF-7 data, for BRCA. The leading or lagging strand was assigned to the TCN motifs as in (Seplyarskiy et al. 2016).

To estimate the statistical significance of the observation that the lagging/leading strand ratio of the APOBEC mutational density is maximal at middle-replicating regions, we repeatedly shuffled mutations between lagging and leading strand for each replication timing bin. For each shuffle, we applied quadratic regression to fit a parabolic curve and to obtain the coefficient of the quadratic term reflecting the curve curvature.

### Gene expression analysis

Gene annotations including gene direction (to infer the sense/antisense strand) were taken from RefSeq (Pruitt et al. 2014). Gene-level transcript abundances quantified by RSEM (Li and Dewey 2011) were downloaded from the Broad TCGA GDC (BRCA 2016; BLCA 2016; HNSC 2016; LUAD 2016; LUSC 2016); estimated gene expression levels in the “scaled_estimate” column, representing TPM values according to the description in TCGA wiki, were used. The values of gene expression were divided into seven intervals (Table S4). Samples with less than six hundred mutations in genes were excluded. Mutational densities in the expression bins were calculated similarly to the densities in replication timing bins, as described above.

### Mutation clusters and model of mutagenesis

Mutation clusters were defined as described previously (Roberts et al., 2013). Briefly, all groups of at least two mutations in which neighboring changes were separated by 10kb or less were identified and the *p*-value for each group was calculated under the assumption that all mutations were distributed randomly across the genome as described previously (Roberts et al. 2012). Groups of mutations were identified as clusters if the calculated *p*-value was than 10^−4^ or less. We also introduced additional strict rules for the analysis of mutational clusters — a particular cluster was considered as an APOBEC-induced cluster if all constituent SBS conformed to the APOBEC signature. Similarly, a mutation cluster was considered as non-APOBEC-induced if all cluster’ SBS did not conform to the APOBEC signature. The regression model of the ABOBEC mutagenesis was defined as *NRD*_*APOBEC*_ (*r, t*) = *β*_0_ + *β*_1_*r* + *β*_2_*t* + *ε*, where *NRD*_*APOBEC*_ is the normalized relative density of APOBEC-induced mutations *NRD*_*APOBEC*_ (*r, t*) = *RD*_*APOBEC*_ (*r, t*)/ ∑_*r,t*_ *RD*(*r, t*). This value is normalized on the sample mutation load to compensate for different time of exposure to mutagens in different samples; *r* is the replication timing, *t* is the gene expression level, *β*_*i*_ are the model coefficients, *ε* is the random error.

## Data Availability Statement

Source codes are available at https://github.com/intersystems-ru/apobec_expression and https://github.com/mkazanov/molcompbio.

## Author Contributions Statement

M.D.K. conceived the study. A.C., B.F., A.K., E.S., J.P. performed the calculations. G.V.P. wrote the code for calculations on the computation cluster. D.V. provided support for performing calculations on the computation cluster. All author contributed to data analysis. M.S.G. and M.D.K. wrote the paper.

## Acknowledgments

We thank Dmitry Gordenin for inspiring us to investigate a role of transcription in the APOBEC mutagenesis, thoughtful discussions and critical reading of the manuscript. This study was partially supported by RFBR via grant 18-29-13011 to M.S.G. and by InterSystems via Innovations Program grant to M.D.K..

## Supplemental Figures

**Figure S1.**
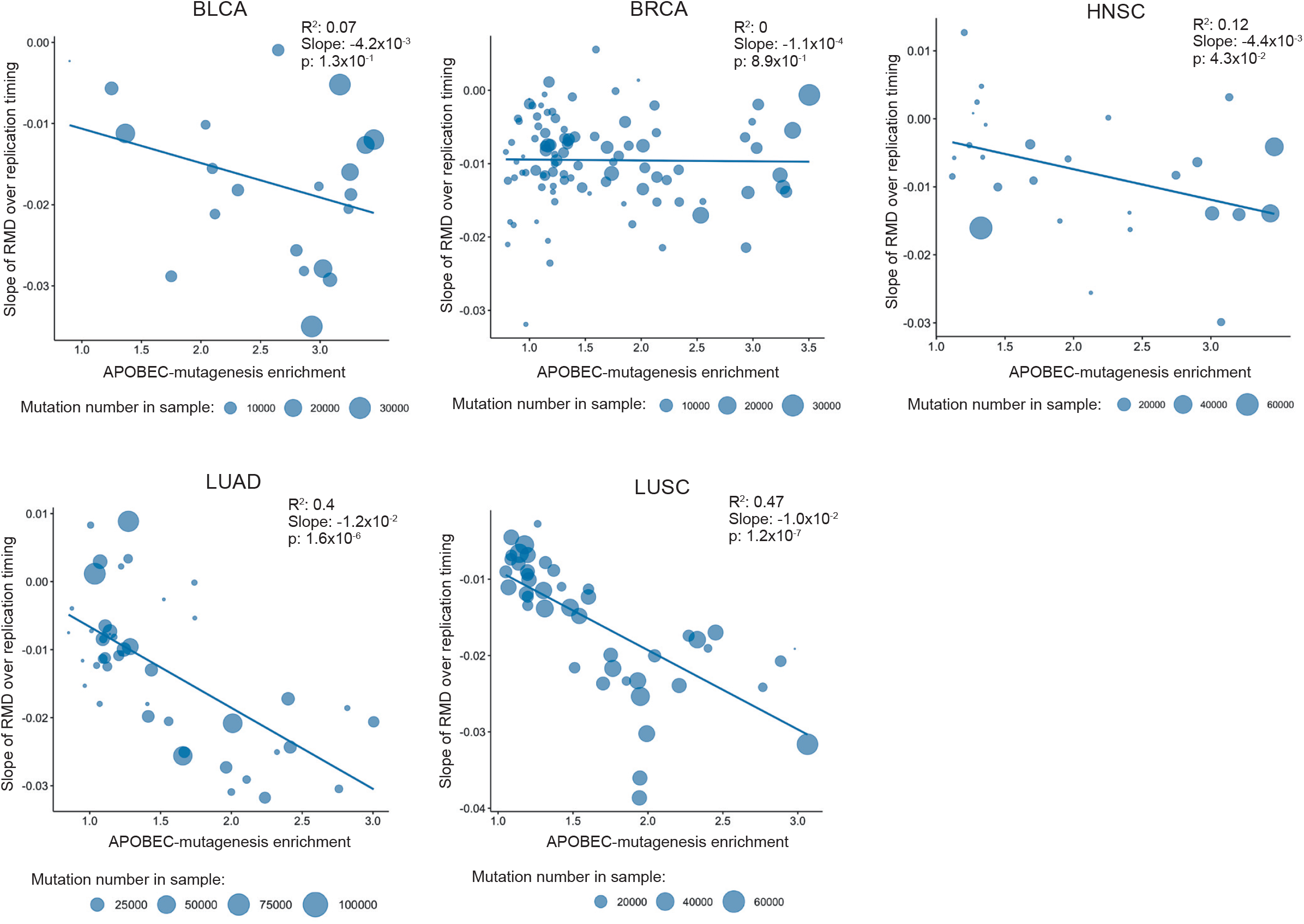
The slopes of the relative mutational density (RMD) distribution (see Methods) of APOBEC-induced SBS (TCW mutational signature) over replication timing as dependent on the activity of APOBEC mutagenesis for samples from five cancer types.

**Figure S2.**
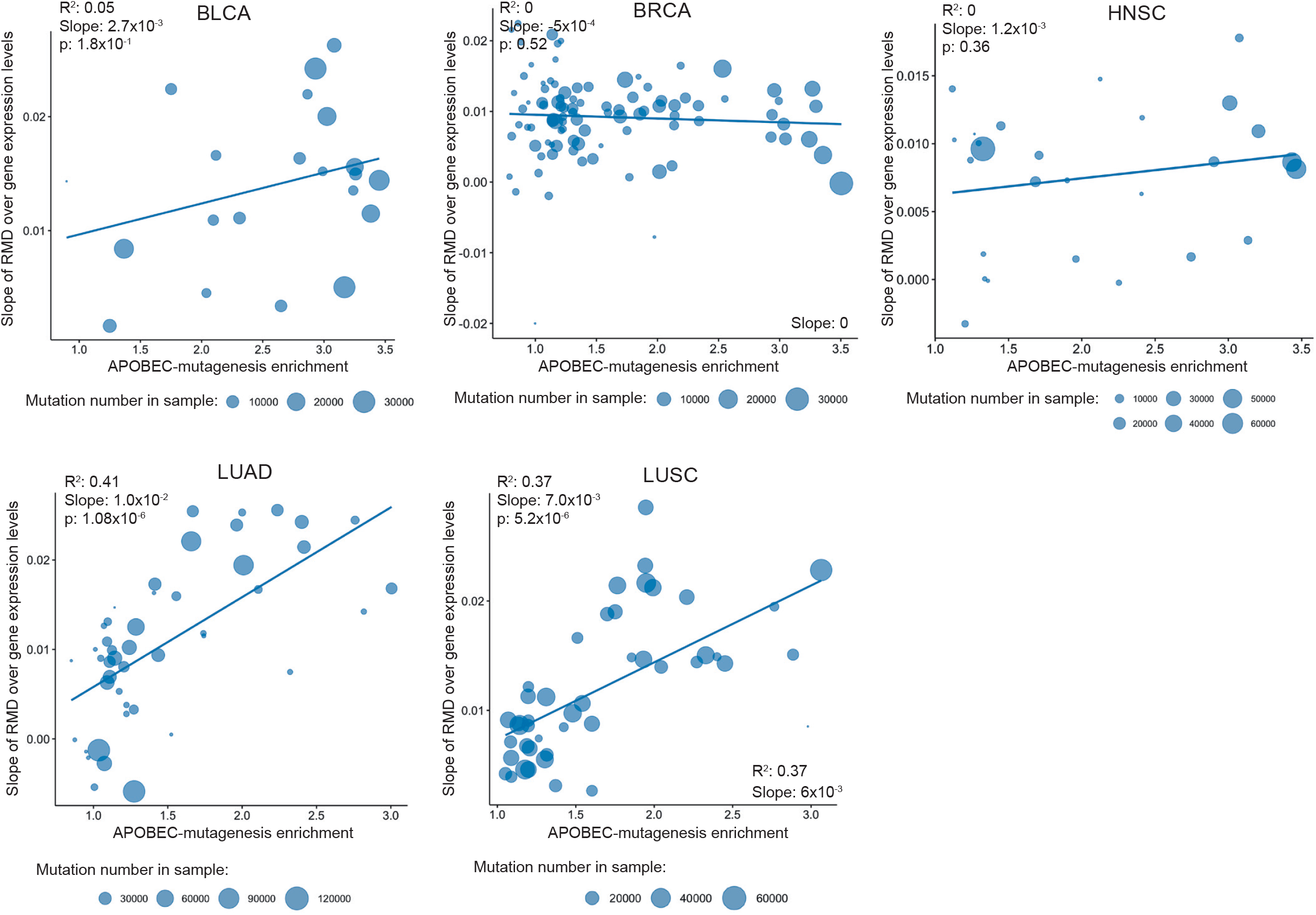
The slopes of the relative mutational density (RMD) distribution of APOBEC-induced SBSs (TCW motif) over gene expression levels as dependent on the activity of APOBEC mutagenesis for samples from five cancer types.

**Figure S3.**
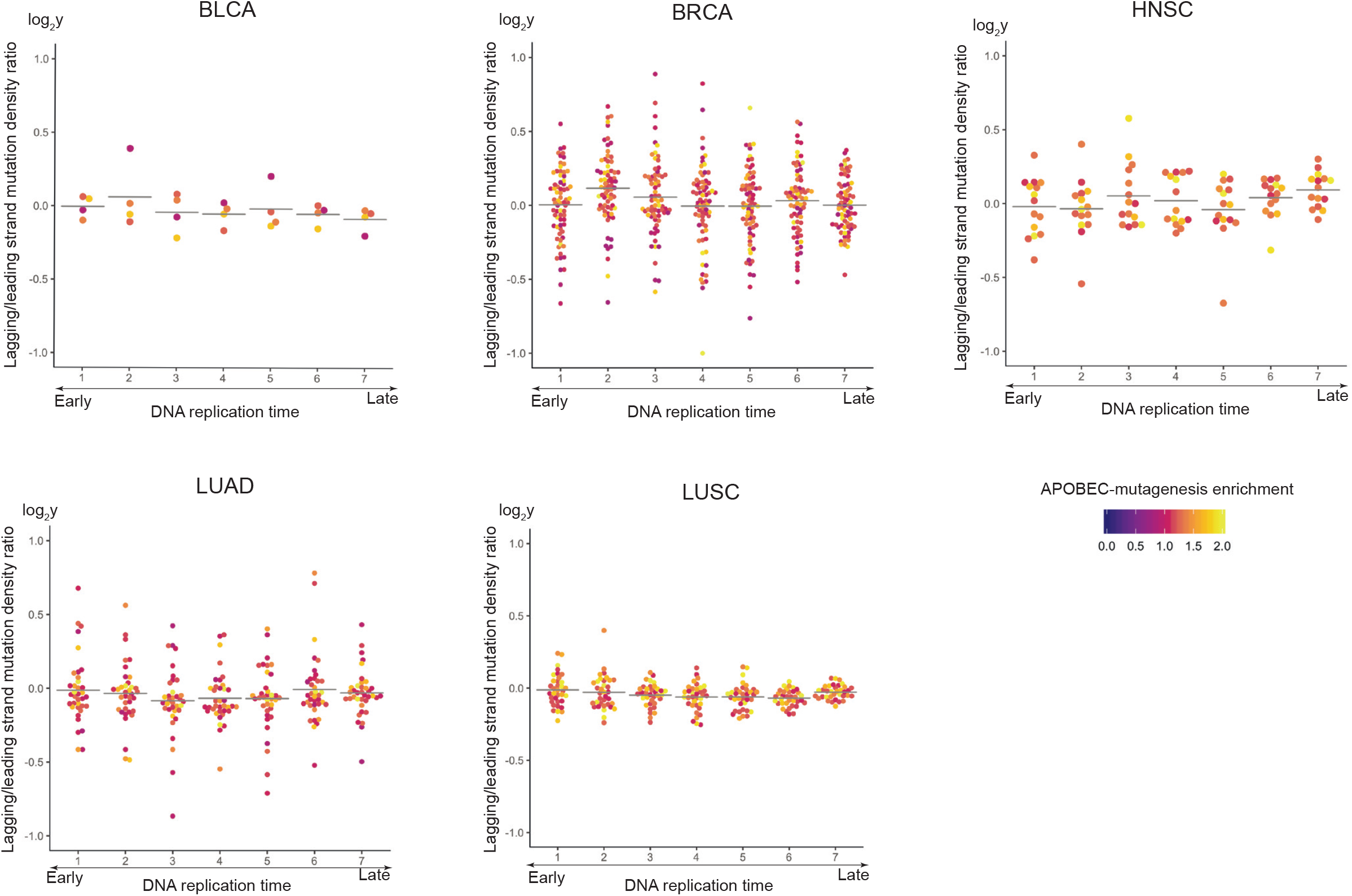
Dependence of the lagging/leading strand ratio of SBS density in cytosines, excluding APOBEC-induced SBS, on the replication timing.

**Figure S4.**
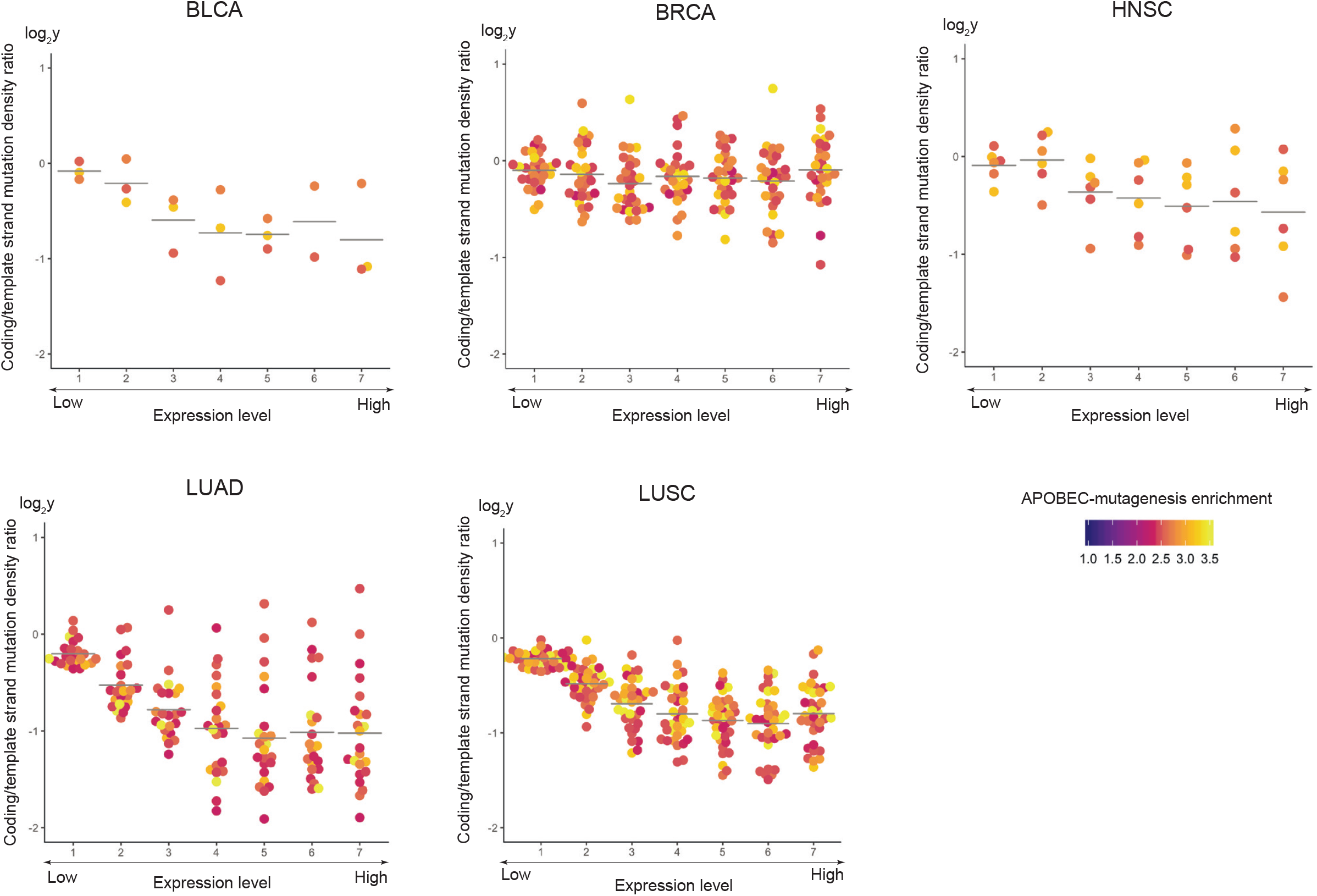
Dependence of the sense/antisense strand ratio of SBS density in cytosines, excluding APOBEC-induced SBS density on the gene expression level.

**Figure S5.**
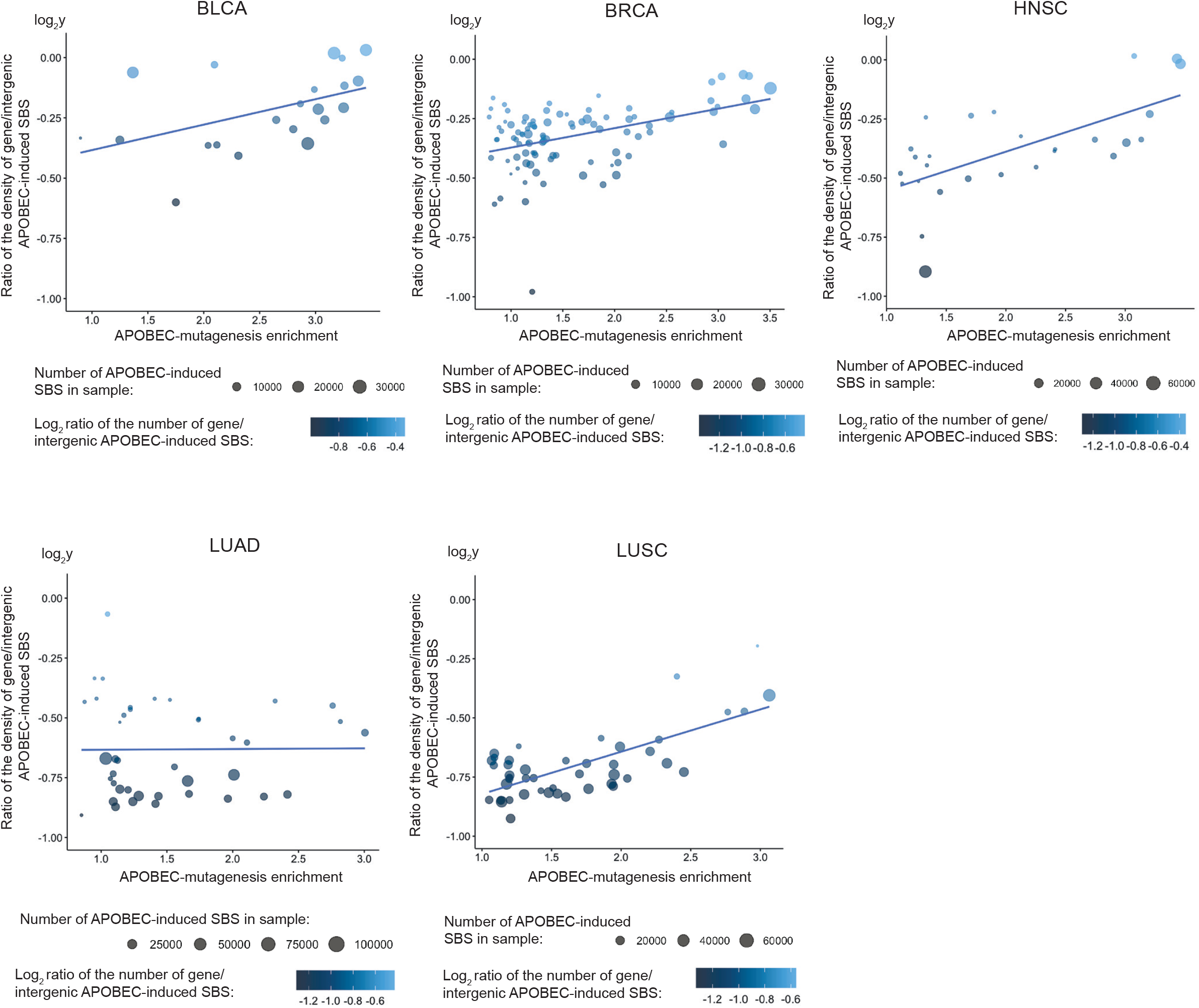
Dependence of the ratio of density and number of APOBEC-induced mutations in gene/intergenic regions on the activity of APOBEC mutagenesis for samples from five cancer types.

**Figure S6.**
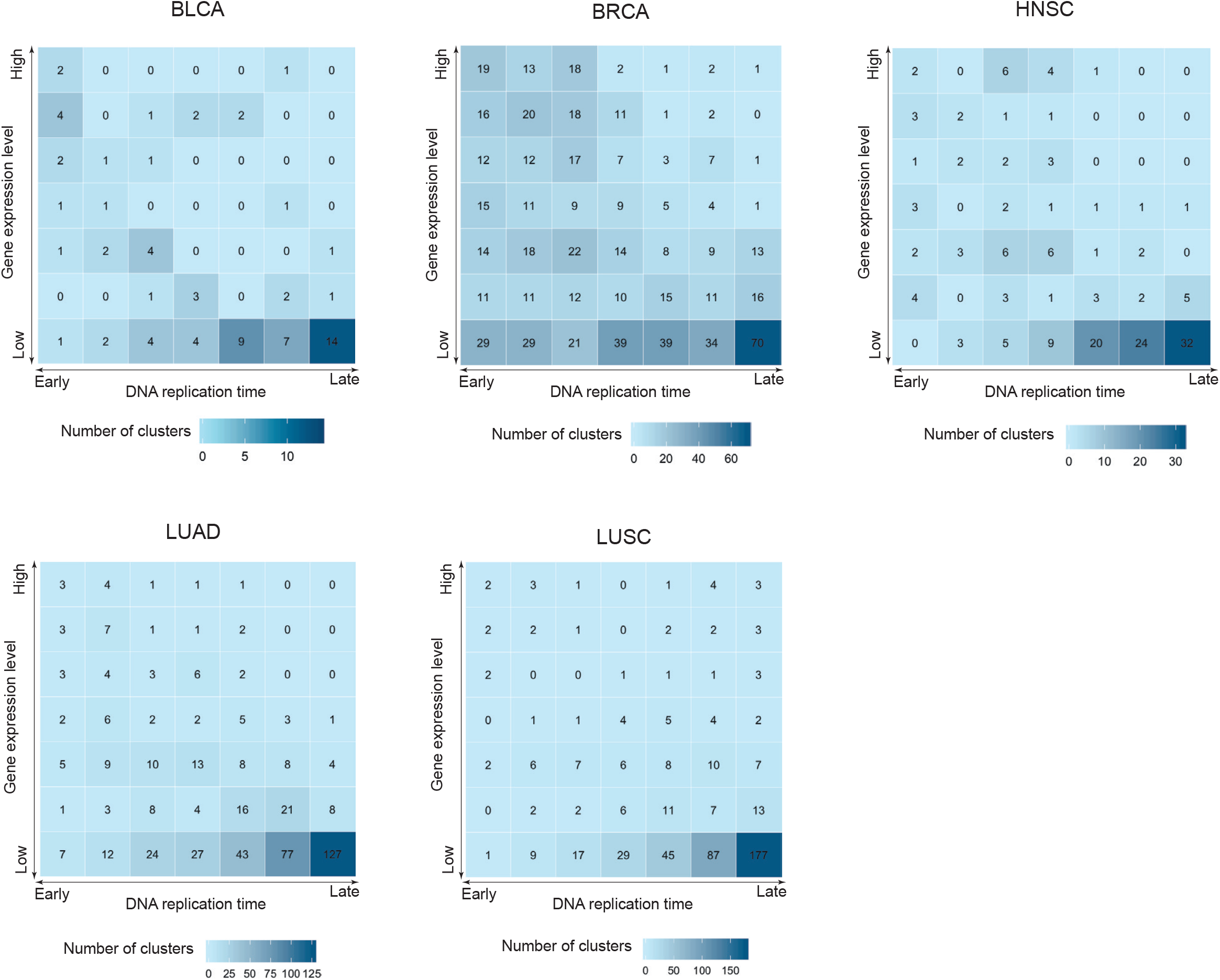
Distribution of non-APOBEC-induced mutation clusters over replication timing and gene expression.

**Figure S7.**
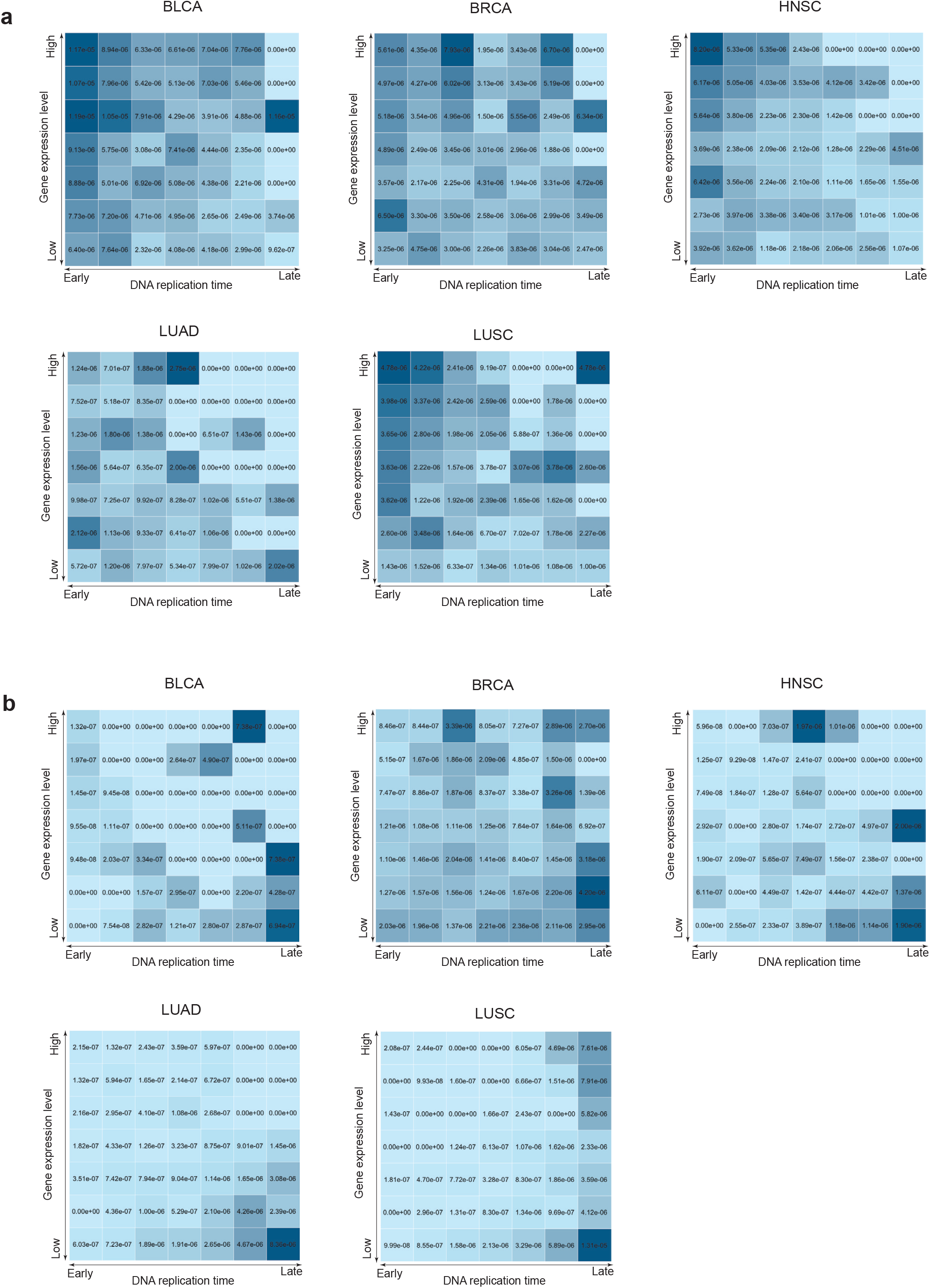
Distribution of the density of (a) APOBEC-and (b) non-APOBEC-induced mutation clusters over replication timing and gene expression.

**Figure S8.**
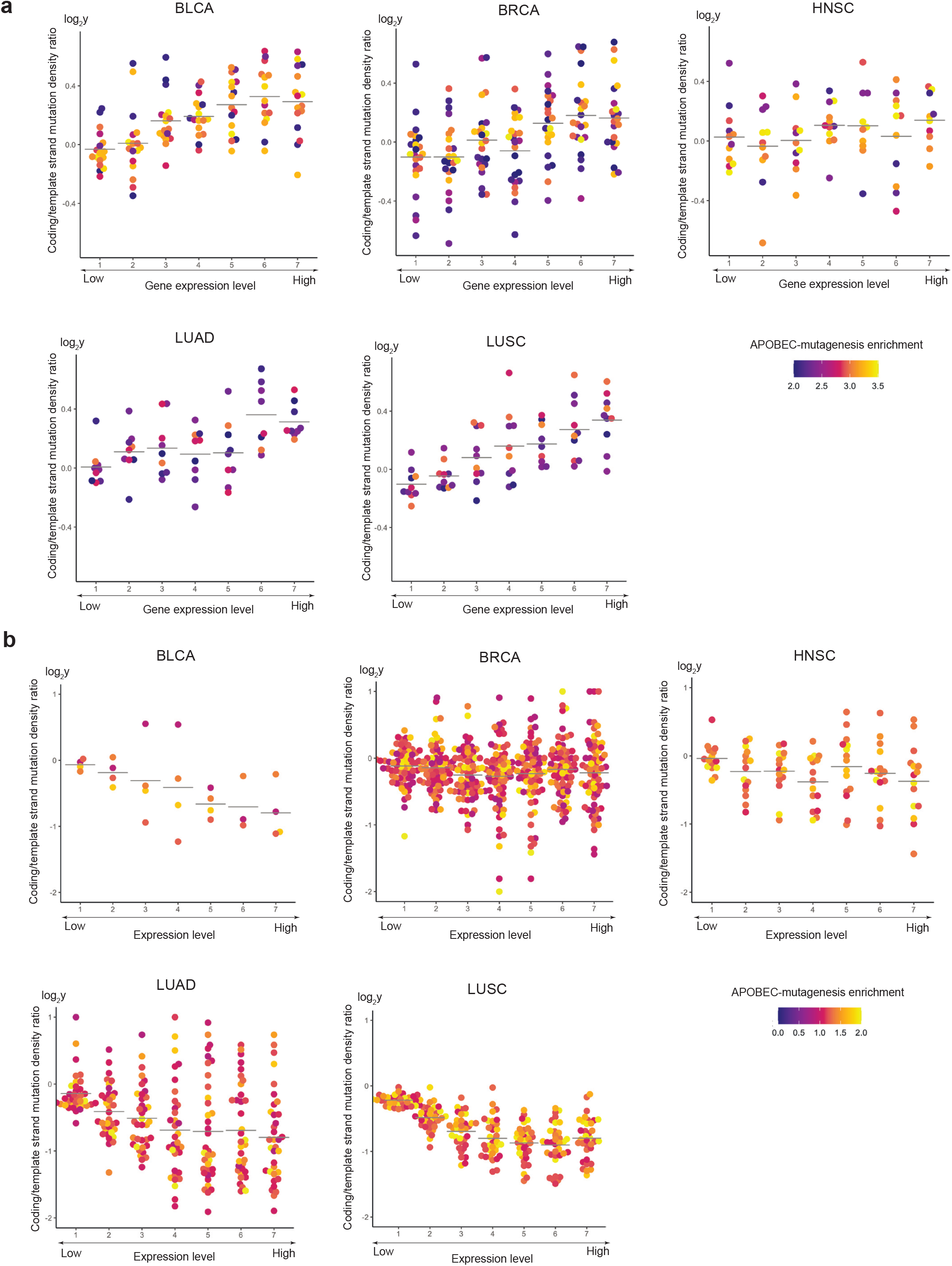
Dependence of the sense/antisense strand ratio of APOBEC-induced SBS density (a) and density of other SBSs in cytosines (b) on the gene expression level. Cancer samples with a low number of mutations and hence higher number of outlier values of sense/antisense strand ratio have not been filtered.

**Figure S9.**
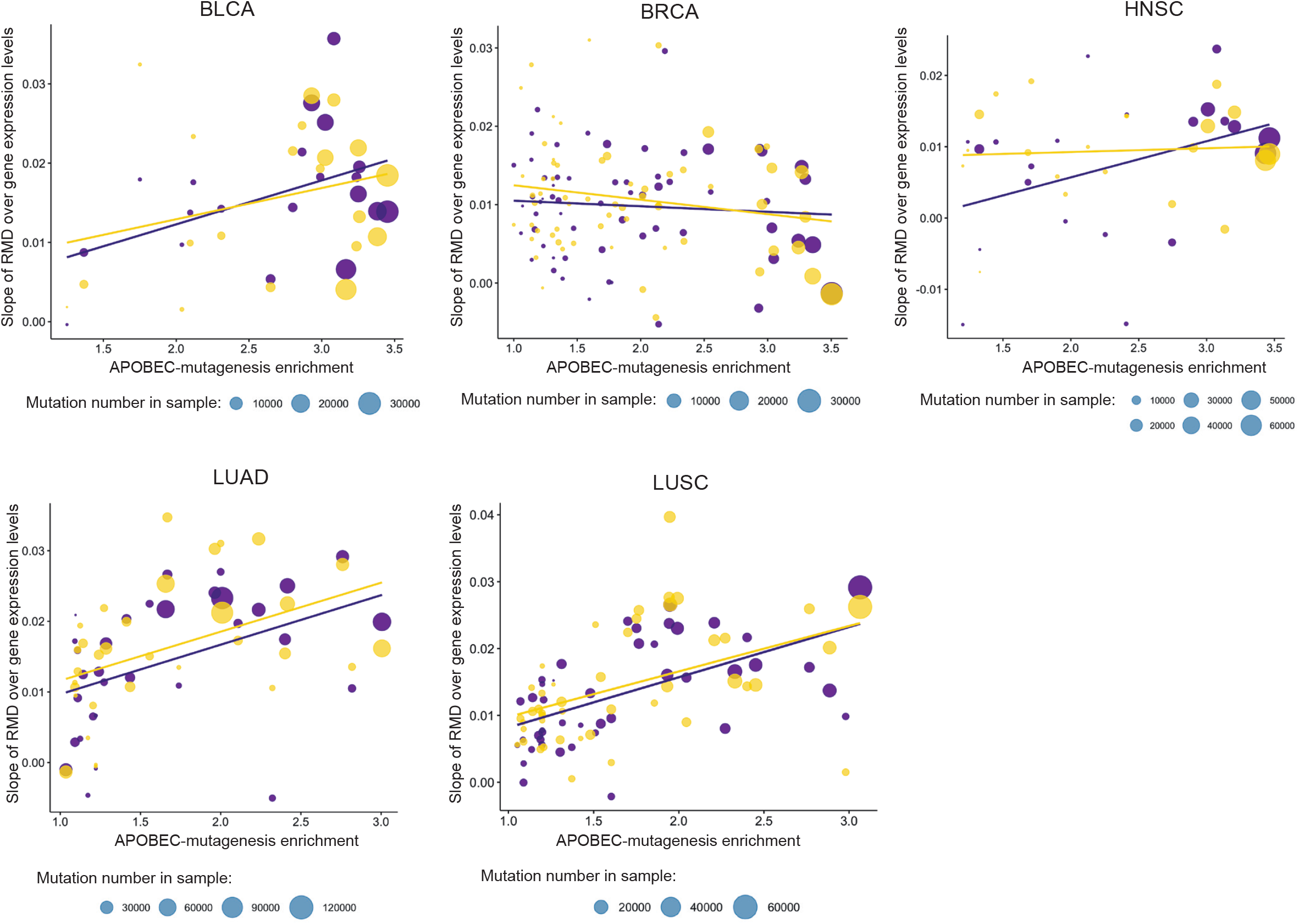
The slopes of the relative mutational density (RMD) distribution of APOBEC-induced SBSs (the TCN motif) over gene expression levels as dependent on the activity of APOBEC mutagenesis for genes transcribed in the direction of replication (violet) and in the opposite direction (yellow).

**Figure S10.**
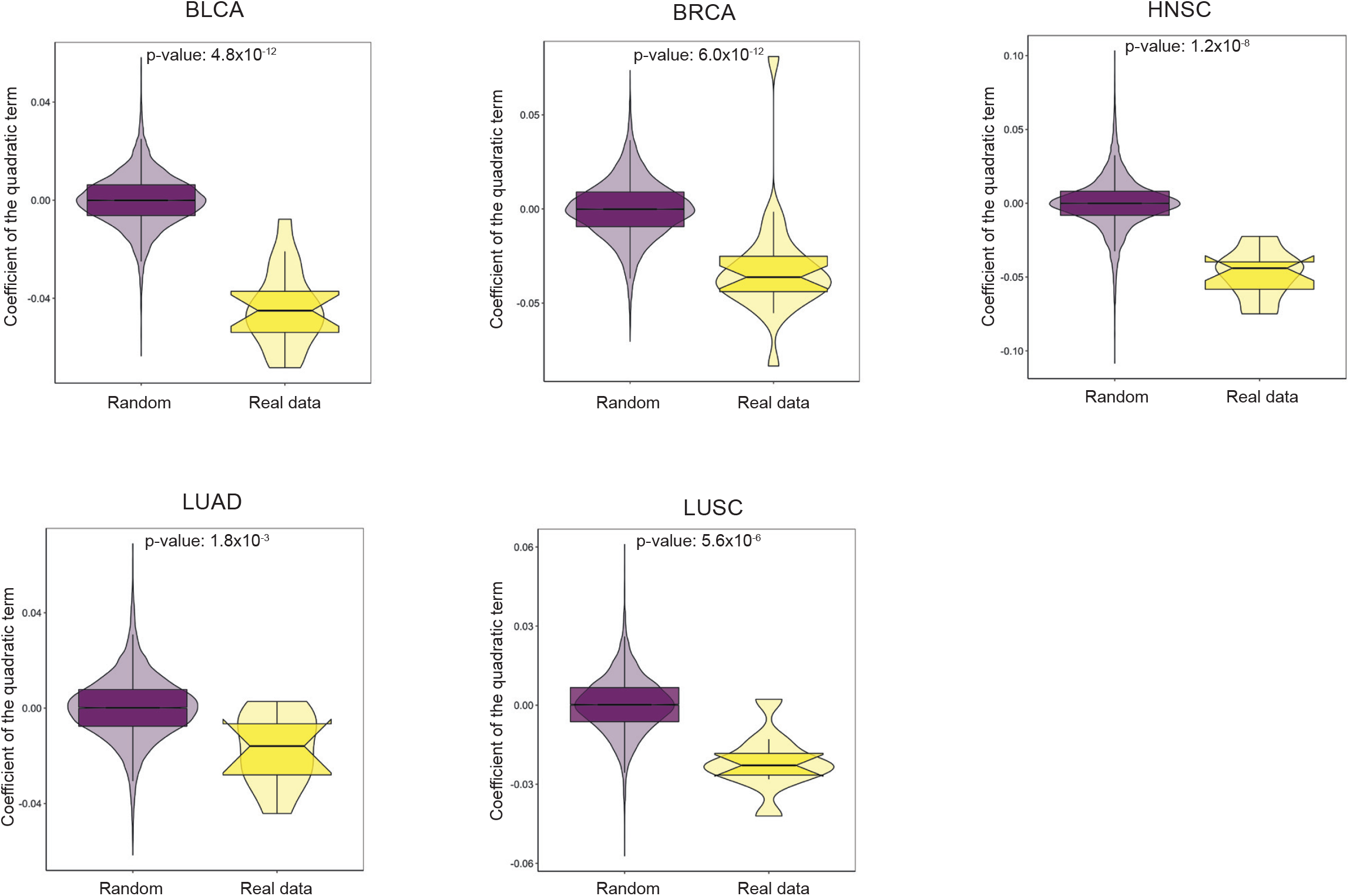
Estimation of the statistical significance for the effect of the highest lagging/leading strand ratio of the APOBEC-induced mutational density at the middle of the replication timing. The distribution of the specified lagging/leading strand ratio along the replication timing was fitted by a quadratic regression and the coefficient of the quadratic term was estimated for each cancer sample. The distribution of the obtained coefficients was compared with the similar distributions obtained from the mutational data where the replication strand was randomly assigned for each mutation.

**Figure S11.**
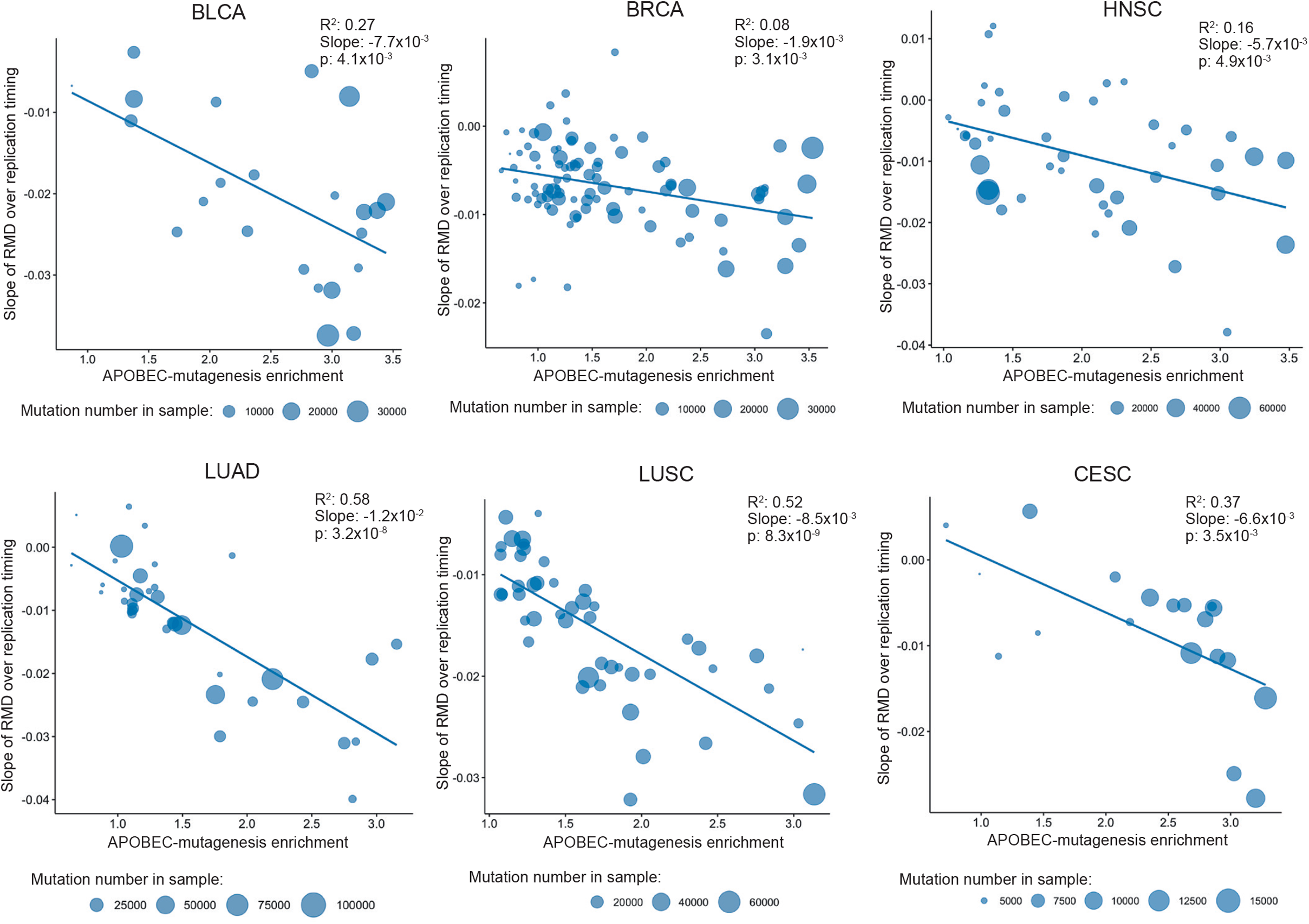
The slopes of the relative mutational density (RMD) distribution of APOBEC-induced SBSs (the TCN motif) over replication timing as dependent on the activity of APOBEC mutagenesis for cancer samples from the PCAWG dataset.

**Figure S12.**
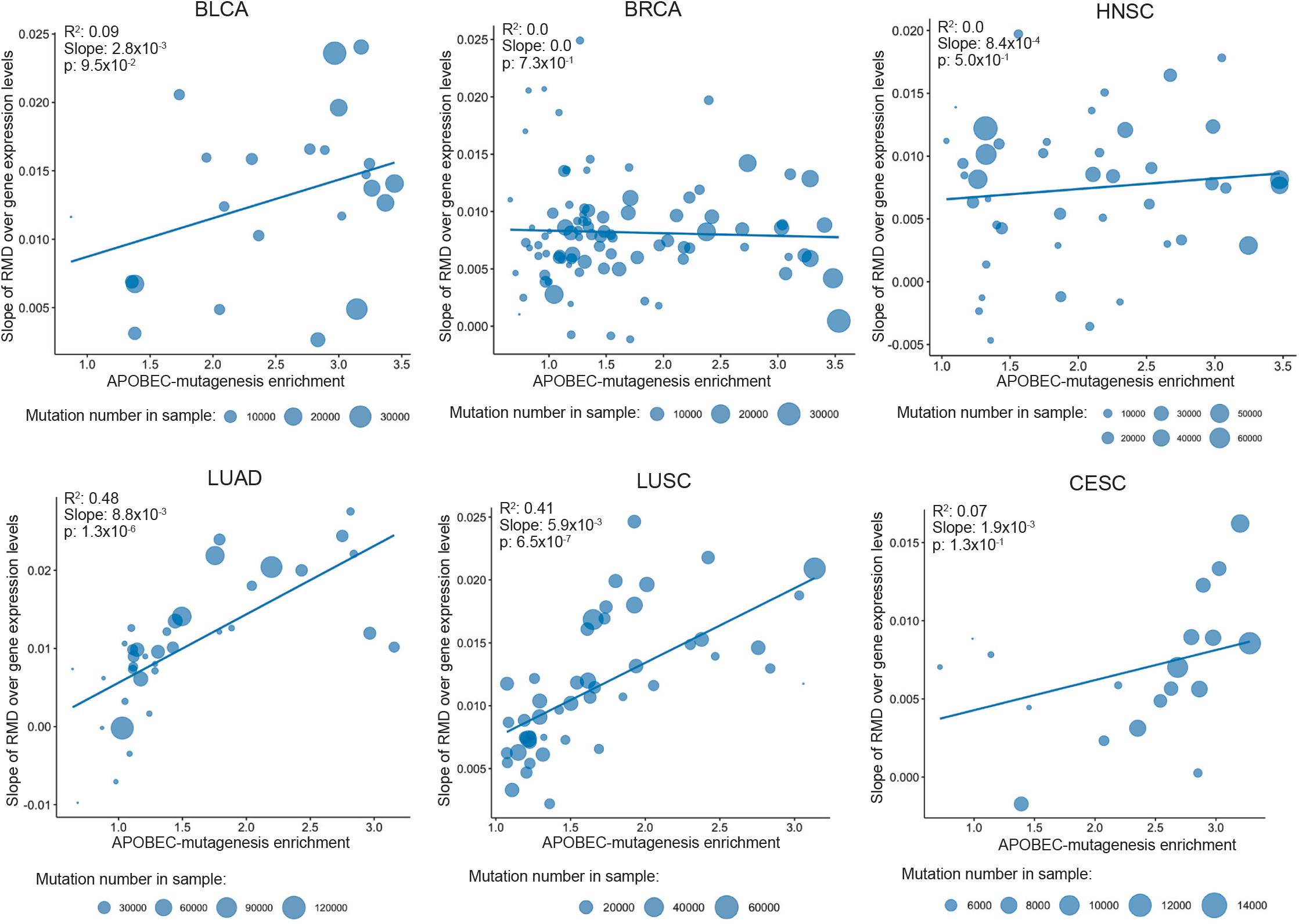
The slopes of the relative mutational density (RMD) distribution of APOBEC-induced SBSs (TCN motif) over gene expression levels as dependent on the activity of APOBEC mutagenesis for cancer samples from the PCAWG dataset.

**Figure S13.**
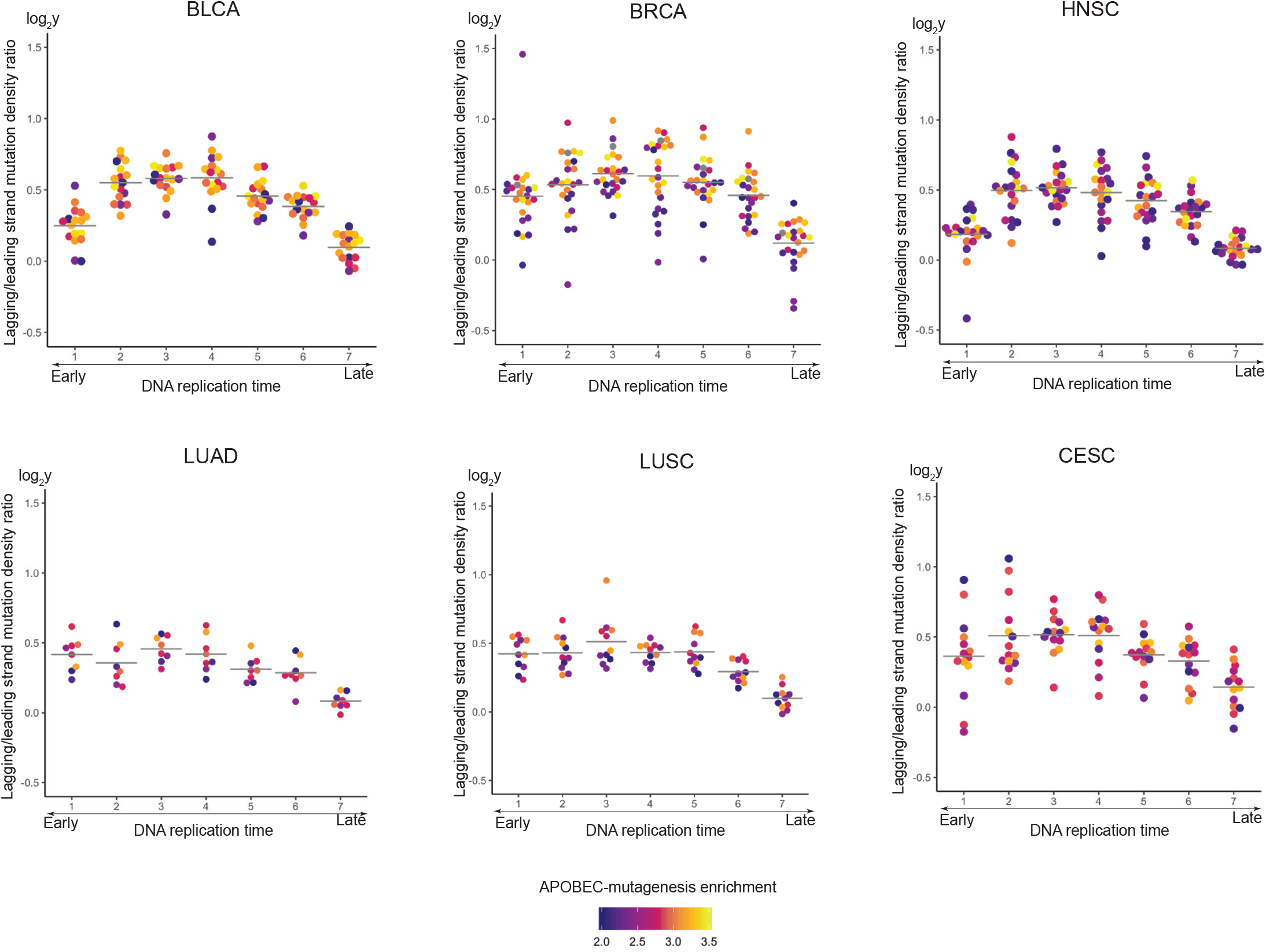
Dependence of the lagging/leading strand ratio of APOBEC-induced SBS density on the replication timing for cancer samples from the PCAWG dataset. The horizontal lines show the mean lagging/leading strand ratio values.

**Figure S14.**
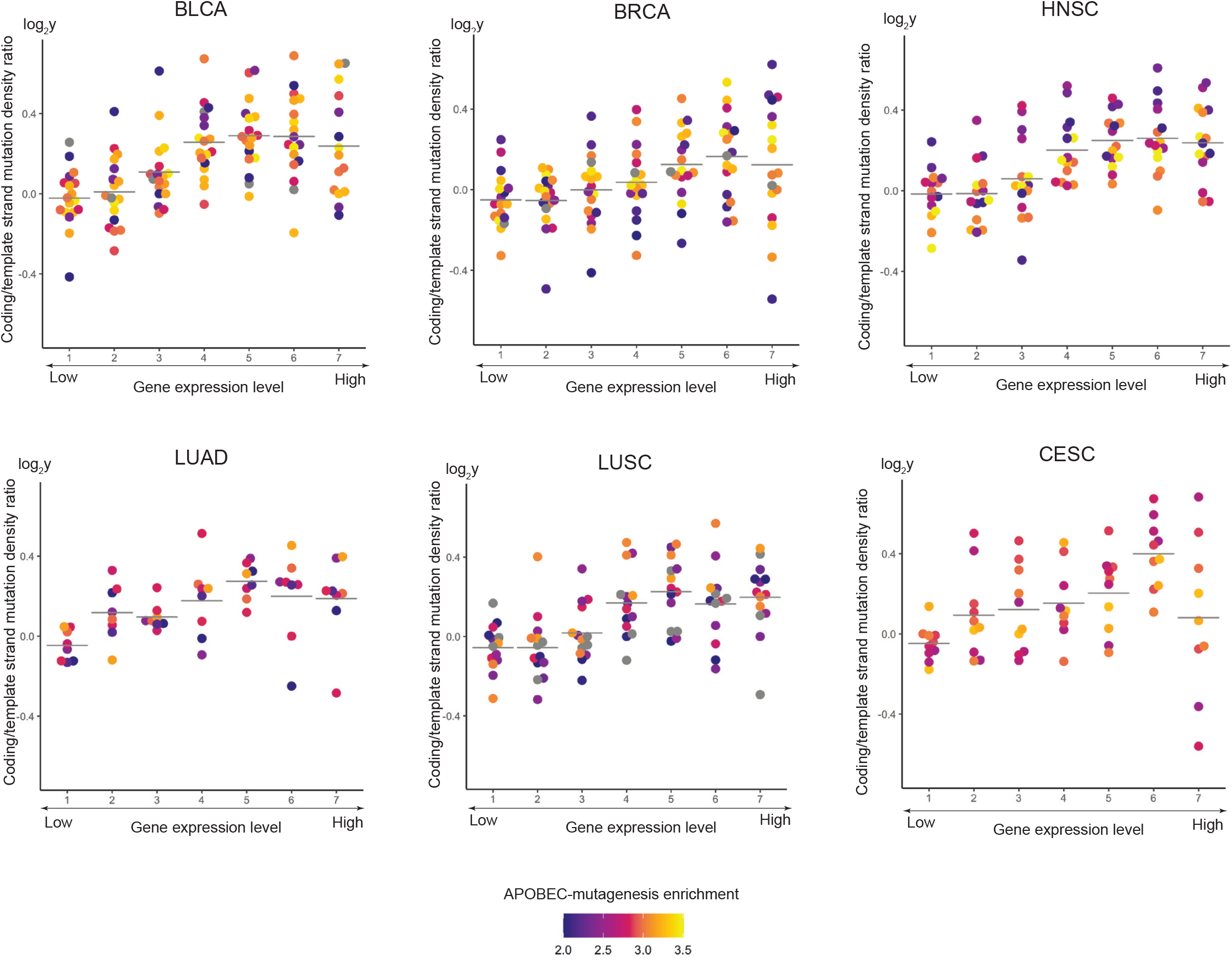
Dependence of the sense/antisense strand ratio of APOBEC-induced SBS density on the gene expression level for cancer samples from the PCAWG dataset. The horizontal lines show the mean sense/antisense strand ratio values.

**Figure S15.**
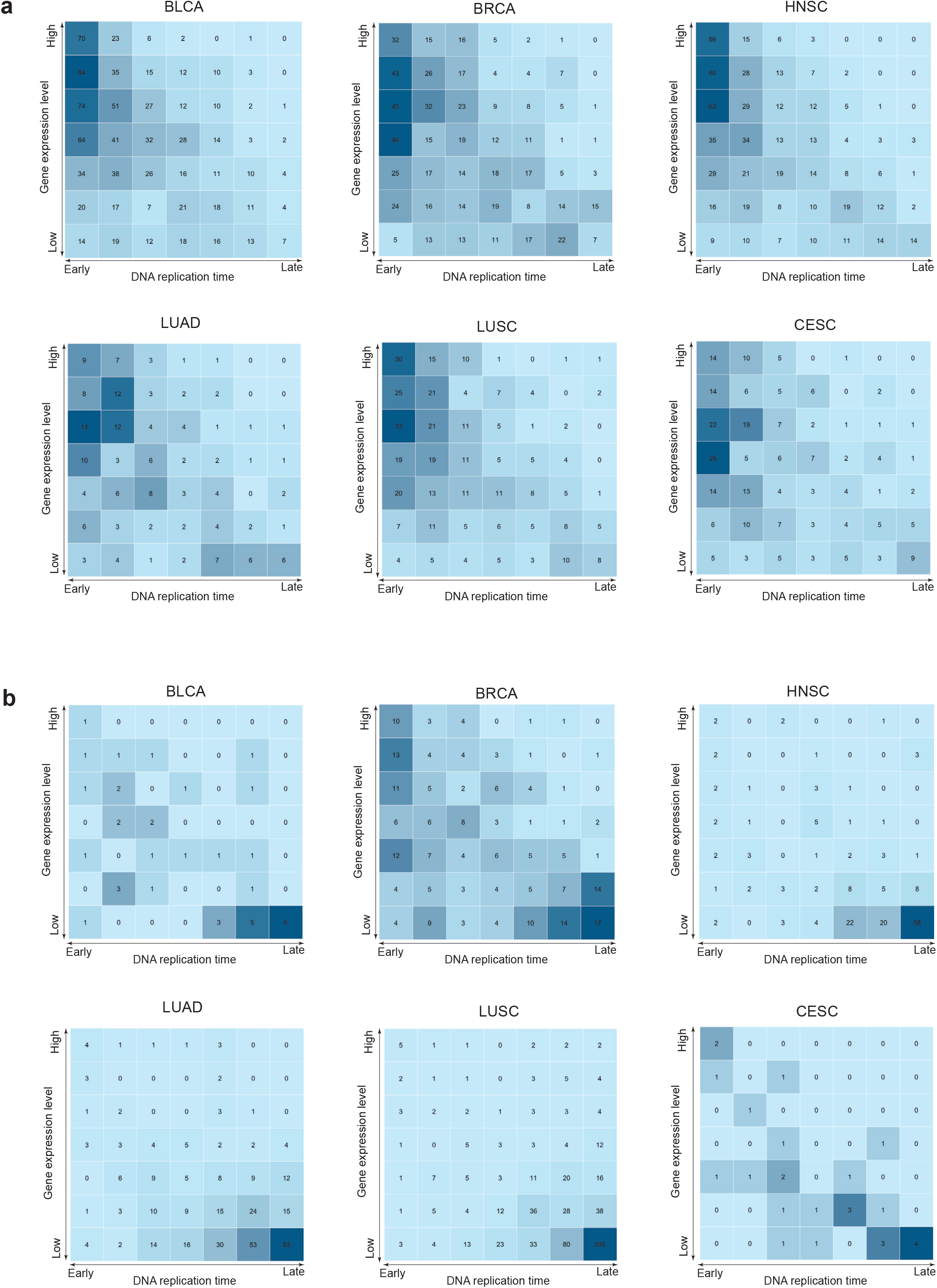
Distribution of the density of (a) APOBEC- and (b) non-APOBEC-induced mutation clusters over replication timing and gene expression for cancer samples from the PCAWG dataset.

**Figure S16.**
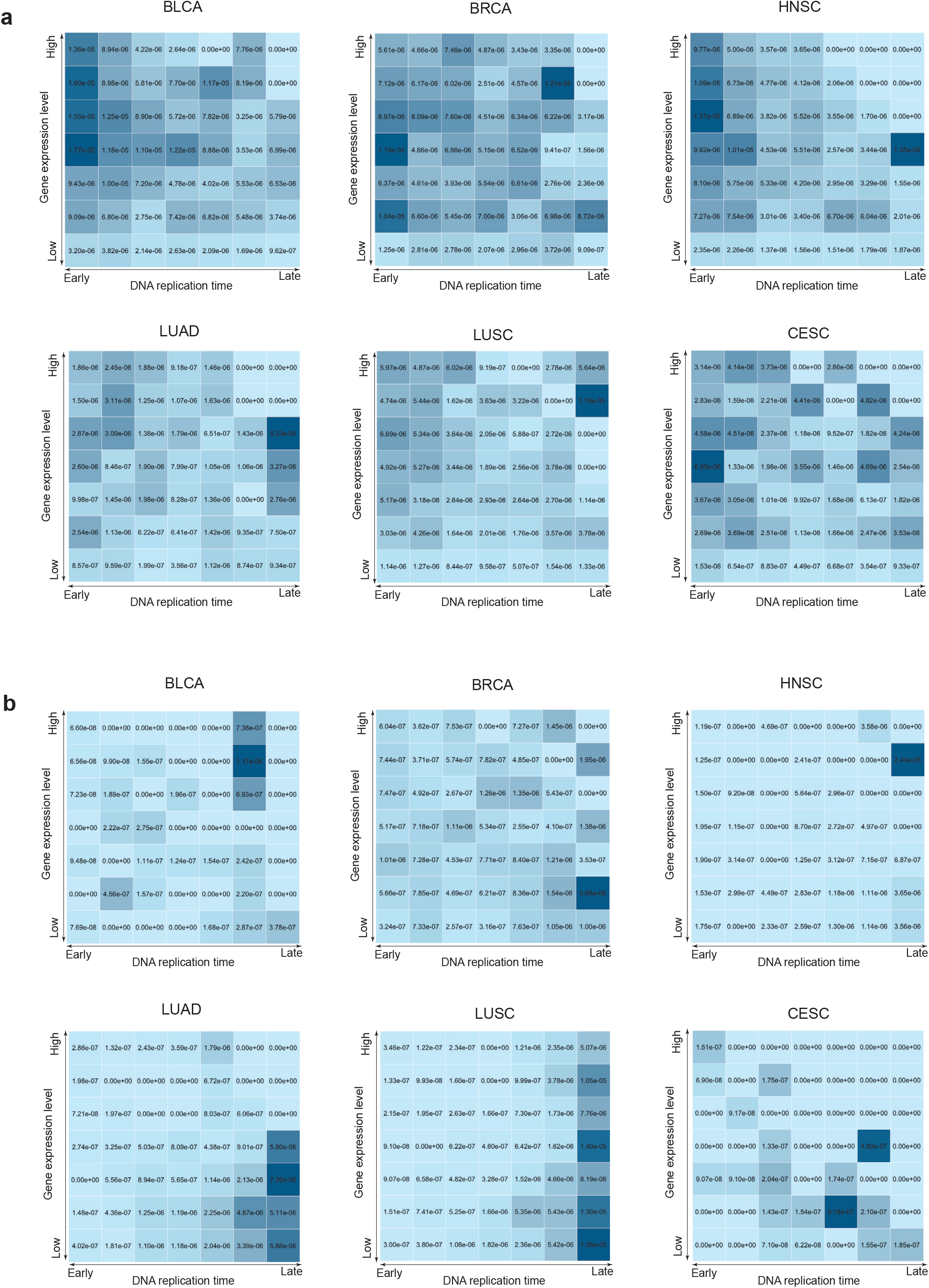
Distribution of the number of (a) APOBEC- and (b) non-APOBEC-induced mutation clusters over replication timing and gene expression for cancer samples from the PCAWG dataset.

**Figure S17.**
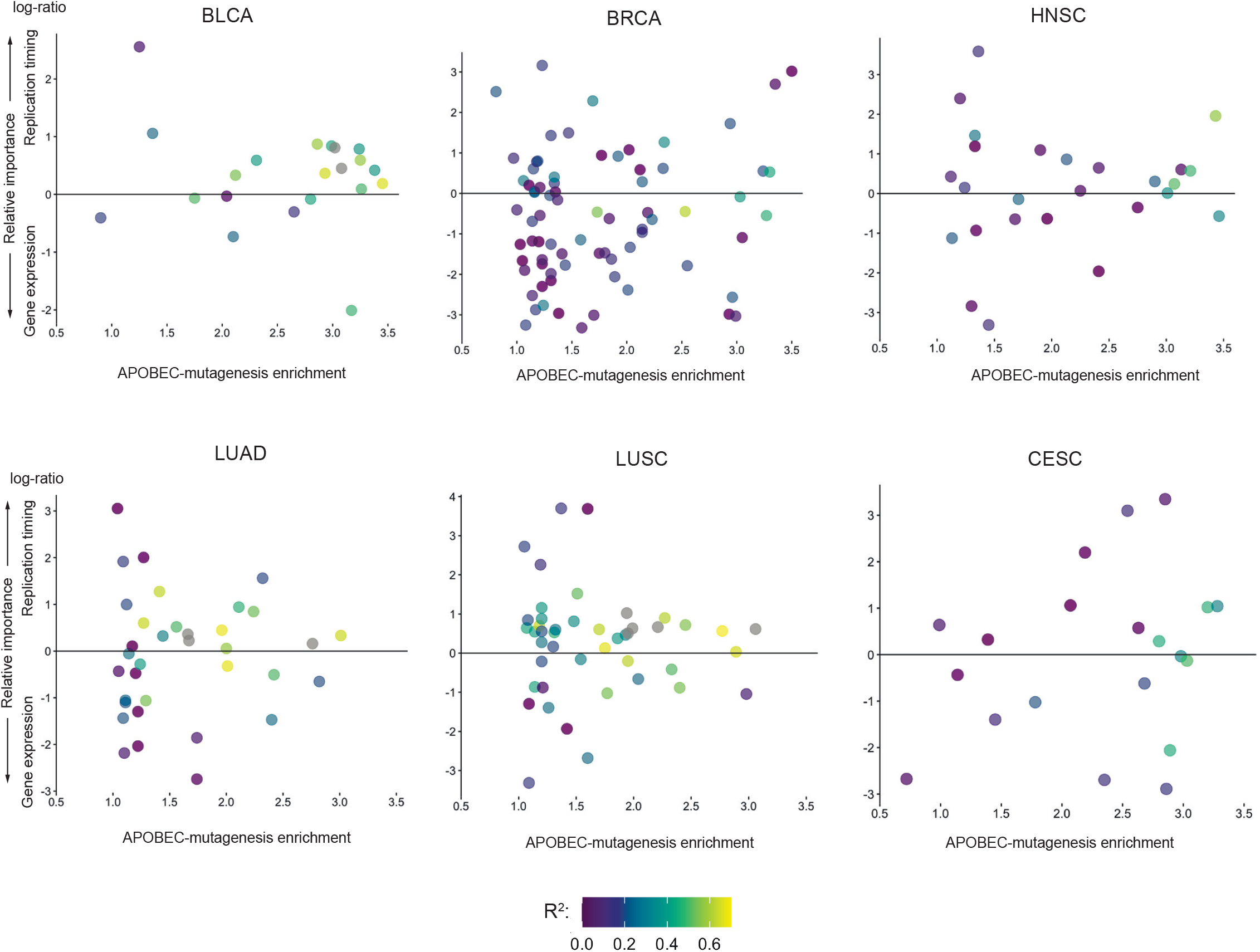
Relative impact of the replication timing and the level of gene expression to the APOBEC-mutagenesis estimated by the LMG method applied to the linear model. The relative importance of the replication timing (I_RT_) and the level of gene expression (I_GE_) was estimated as the increase in R^2^ averaged over different regressor orderings. The vertical axis represents the logarithm of the ratio of the relative importance of replication timing to the relative importance of gene expression, i.e. log(I_RT_/I_GE_).

**Figure S18.**
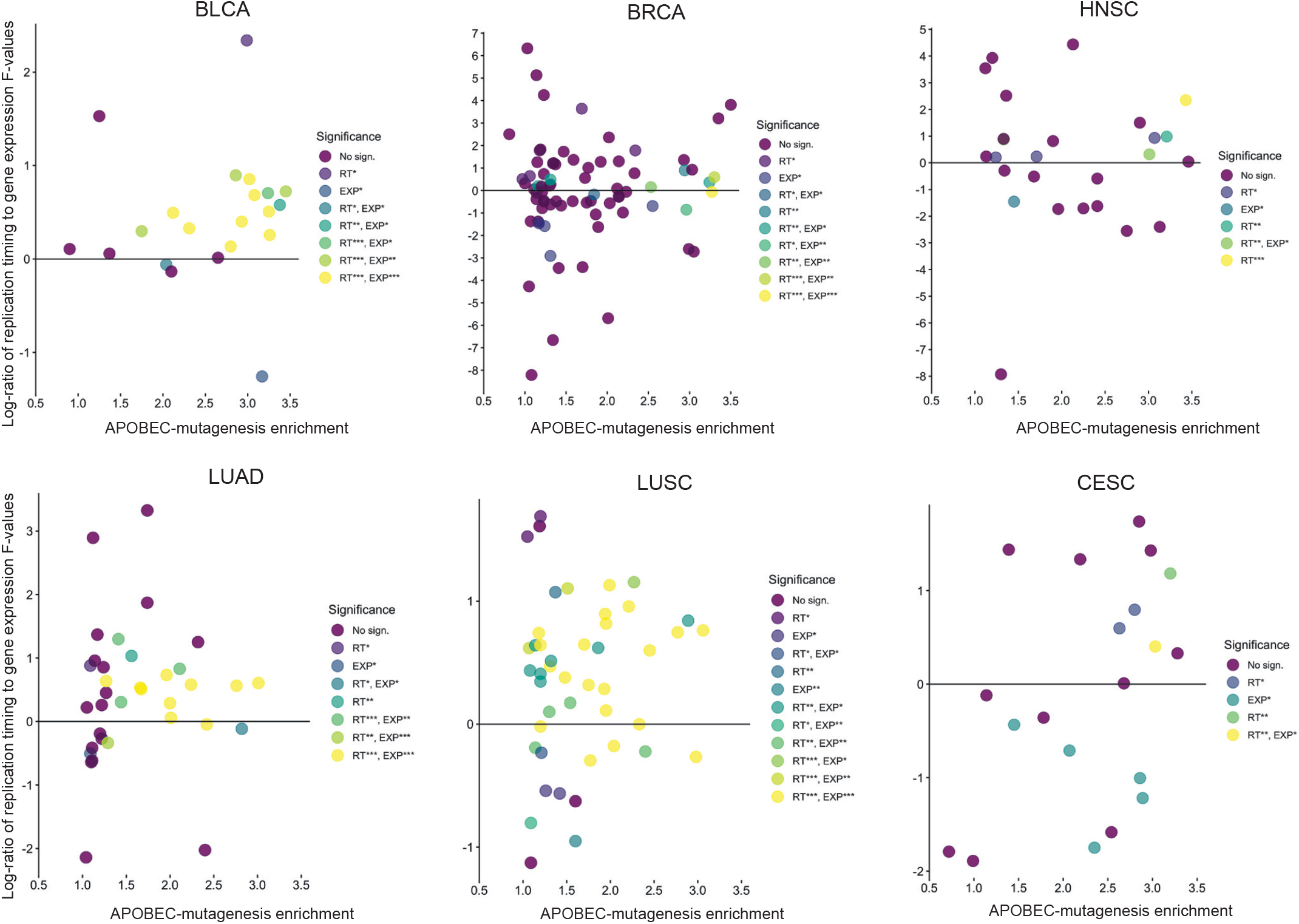
Comparison of the influence of the replication timing and the level of gene expression on APOBEC-mutagenesis, as estimated by the two-way Analysis of Variance (ANOVA). The relative importance of the first (replication timing) and second (level of gene expression) factors is presented as the log-ratio of calculated F-values. Statistical significances of factors are visualized as RT*** (p<0.001), RT** (p<0.01) and RT* (p<0.05) for the replication timing, and as EXP*** (p<0.001), EXP** (p<0.01), EXP* (p<0.05) for the level of gene expression.

**Table S1.** Replication timing bins for IMR90 cell

| Replication bin | Minimal replication timing value | Maximal replication timing value |
| --- | --- | --- |
| 1 | 75 | 90 |
| 2 | 64 | 75 |
| 3 | 52 | 64 |
| 4 | 40 | 52 |
| 5 | 28 | 40 |
| 6 | 13 | 28 |
| 7 |  | 13 |

**Table S2.** Replication timing bins for NHEK cell

| Replication bin | Minimal replication timing value | Maximal replication timing value |
| --- | --- | --- |
| 1 | 67 | 80 |
| 2 | 60 | 67 |
| 3 | 51 | 60 |
| 4 | 41 | 51 |
| 5 | 32 | 41 |
| 6 | 22 | 32 |
| 7 |  | 22 |

**Table S3.** Replication timing bins for MCF7 cell

| Replication bin | Minimal replication timing value | Maximal replication timing value |
| --- | --- | --- |
| 1 | 68 | 86 |
| 2 | 60 | 68 |
| 3 | 50 | 60 |
| 4 | 40 | 50 |
| 5 | 30 | 40 |
| 6 | 19 | 30 |
| 7 |  | 19 |

**Table S4.** Gene expression bins

| Expression bin | Min expression value | Max expression value |
| --- | --- | --- |
| 1 | 0 | 25 |
| 2 | 25 | 100 |
| 3 | 100 | 300 |
| 4 | 300 | 550 |
| 5 | 550 | 1000 |
| 6 | 1000 | 2000 |
| 7 | 2000 |  |

## Supplemental Files

Supplemental File 1: https://drive.google.com/file/d/1KRahsYg6BUOG6B4FMgyw0JGrBBRT8SSB/view?usp=sharing

Supplemental File 2: https://drive.google.com/file/d/13LYH-gk1IS5BDKOgb8IoRpuNKn0097OS/view?usp=sharing

Supplemental File 3: https://drive.google.com/file/d/1PxMvAB1qg3OBgrsuGT1uQM8sQy7Or8MT/view?usp=sharing

Supplemental File 4: https://drive.google.com/file/d/1GY7LfFZ2EGjQl4NHg91F9tOXl0TT3xog/view?usp=sharing

